# Fubylation as a Druggable Ubiquitin-Like Conjugation System Controlling Hippo Pathway Activity

**DOI:** 10.1101/2025.07.06.663376

**Authors:** Maya L. Bulos, Edyta M. Grzelak, Kayla Nutsch, N. G. R. Dayan Elshan, Arnab K. Chatterjee, Michael J. Bollong

## Abstract

The functionally and evolutionarily conserved Hippo-YAP signaling pathway plays a critical role in regulating cellular proliferation, organ size control, and regeneration. Accordingly, activators of YAP-driven transcription hold therapeutic promise for treating disease states driven by insufficient proliferative repair, yet only a handful of pharmacological mechanisms exist for augmenting YAP activity. Here we report the discovery of sCMF231, a small molecule activator of YAP discovered from high throughput screening that acts by targeting the poorly characterized ubiquitin-like protein FUBI. Using canonical ubiquitin conjugation machinery—UBA1, UBE2C, and APC/C—FUBI covalently modifies the Hippo pathway protein Annexin A2, reinforcing its YAP suppressive role at the plasma membrane. Binding of sCMF231 to FUBI discourages its conjugation to Annexin A2, resulting in the membrane delocalization of Annexin A2 and a liberated, transcriptionally active form of YAP. This work provides the first definitive evidence of covalent modification of proteins by FUBI, a post-translational modification termed fubylation, and defines how fubylation regulates the activity of a central growth pathway.

## Introduction

The conserved Hippo pathway regulates cellular proliferation, ectopic growth, and stem cell function in animals. Initially discovered from organ overgrowth genetic screens in *Drosophila*, the core of the pathway consists of a linear kinase cascade, relaying a range of environmental signs to the kinases sterile 20-like kinase 1 (MST1) and MST2 and large tumor suppressor kinase 1 (LATS1) and LATS2^1,2^. Hippo activation leads to the subsequent activation and phosphorylation of MST1/2 and then LATS1/2, resulting in the phosphorylation, inactivation, and subsequent cytoplasmic degradation of the downstream transcriptional co-activator Yes-associated protein (YAP)^3–6^. When the Hippo pathway is inactivated, unphosphorylated YAP translocates to the nucleus where it enacts a pro-proliferative and pro-survival transcriptional program through essential interactions with TEA domain transcription factors (TEAD1-4)^7^.

Several other proteins and mechanisms sense and relay information about cellular density, polarity, and stress to the Hippo pathway, acting to restrain YAP transcriptional activation. Among these are scaffolding proteins, such as Salvador homolog 1 (SAV) and MOB kinase activator 1A and 1B (MOB1A/B) which assist MST1/2 in recruiting and phosphorylating LATS1/2^8–11^. Another central scaffolding protein, Neurofibromin-2 (NF2) templates the phosphorylation and activation of LATS1/2 by MST1/2 at the cell membrane^12^. More recently, other proteins have also been shown to be central modulators of Hippo pathway activity, including Annexin A2 (ANXA2), which localizes to adherens and tight junctions^13,14^, shepherding YAP and MST2 together at the cell membrane^15^. While several models have attempted to describe a single mechanism for how Hippo pathway signaling occurs at the cell membrane with known scaffolding components, the discovery of ANXA2 and other similar proteins make it clear that likely many different interaction ‘nodes’ exist at the cell membrane to relay inhibitory phosphorylation signals to YAP. Accordingly, it is of key importance to better define the spectrum of such nodes and delineate their YAP suppressive mechanisms^14^.

Among known pro-regenerative factors in mammals, YAP activation plays a central role in promoting regenerative organ repair by increasing cellular proliferation and survival^16^. For example, in the intestine, upon acute injury, YAP is required to regenerate the mucosal epithelium, expanding populations of transit amplifying cells^17^. Similarly, in the skin, nuclear levels are augmented in keratinocyte progenitor cells to promote cell epidermal regeneration in the post-wound setting^18^. YAP activity is similarly augmented in the liver following partial hepatectomy and has also been found to play a role in regeneration of the corneal epithelium, the healing of bone fractures, among many other tissues^19–21^. YAP activation via genetic engineering stimulates repair in both regenerating and canonically non-regenerating organs like the heart^22,23^. Cardiac regeneration is particularly promising, as YAP activation via genetic inactivation of SAV effectively reverses heart failure in mice and the more translationally relevant pig model, suggesting that small molecules which activate YAP in cardiomyocytes may be a therapeutically viable approach for reversing disease progression in a large swath of the population^24,25^.

Despite the promise YAP holds in broadly promoting regenerative organ repair, pharmacological mechanisms to activate YAP are understudied, and there are no approved YAP activating drugs. Despite this, examples from our lab and others have indicated that small molecule stimulators of YAP activity show impressive efficacy in animal models of disease^26^. Efforts from others have principally focused on inhibiting the Hippo pathway kinases, MST1/2 and LATS1/2^27,28^. However, these kinases are involved in numerous cellular processes apart from their roles in the Hippo pathway, including cytokinesis, cell motility, transcription, and immune responses, which may lead to undesirable phenotypes in the context of a pro-proliferative agent^27,29–33^. Accordingly, we posit that a superior approach would involve drugging Hippo signaling in a contextually selective manner, impeding relay of phosphorylation to YAP when the pathway is activated but not by inhibiting Hippo kinases in their active sites.

To better drug and to study Hippo suppressive mechanisms, we have performed several historical reporter-based screens of large chemical libraries to identify new YAP activating pharmacological tools. From these efforts, we describe here the discovery of a small molecule that robustly and selectively activates YAP-driven transcription. We define this compound as binding to the poorly studied ubiquitin-like protein FUBI, inhibiting its capacity to functionally conjugate to and facilitate the membrane localization of ANXA2. We further characterize writers of fubylation, known ubiquitin conjugation machinery, and provide the first definitive proof of fubylation on a protein substrate.

## Results

### Identification of sCMF231 as a YAP activator

We previously established a screening platform using a HEK293A cell line possessing a stably integrated cassette of eight copies of the TEAD-binding element upstream of luciferase (293A-TEAD-LUC) from which we identified several novel YAP-activating small molecules. Among chemical scaffolds identified, one quinazoline containing compound, sCGM824, was found to activate luciferase activity in 293A-TEAD-LUC cells with a half maximal efficacious concentration (EC_50_) of 1.6 µM (Fig. S1A). A structure activity relationship (SAR) study from 24 commercially available analogs containing the 4-quinazolamine core of sCGM824 identified sCMF231, which dose-dependently induced TEAD-LUC reporter activity with an EC_50_ of 1.05 µM and displayed no toxicity at tested concentrations (≤20 µM; Fig. 1A, B, Fig. S1B, C). sCMF231 was found to significantly increase the levels of canonically YAP-controlled transcripts *ANKRD1, CYR61,* and *CTGF* in HEK293A cells, as assessed by RT-qPCR (Fig. 1C). Furthermore, Western blotting revealed that sCMF231 treatment concentration-dependently decreases the phosphorylation of YAP in HEK293A cells within 6 hours (Fig. 1D). To further confirm sCMF231 activates a YAP-driven transcriptional program, we performed RNA-sequencing (RNA-seq)-based expression profiling of HEK293A cells, finding that sCMF231 (10 µM) robustly and selectively increased annotated YAP transcriptional targets, as determined by differential expression analysis (DESeq) and gene set enrichment analysis (GSEA)^7,34^ (Fig. 1E, F). We additionally knocked down YAP transcript using a previously reported lentiviral shRNA and found that 10 µM treatment with sCMF231 failed to increase *ANKRD1, CYR61,* and *CTGF* transcript levels in this context, suggesting the compound is dependent on YAP for activating the observed transcriptional profile (Fig. S1D). We also showed that 5 µM sCMF231 treatment promoted the clonal expansion of immortalized human keratinocytes (HaCaT), as indicated by rhodamine B staining and increased nuclei counts after 7 days (Fig. S2A, B), a result consistent with the pro-proliferative effects observed with other pharmacological YAP activators^15,35^. Interestingly, co-treatment of sCMF231 and PY-60, an ANXA2-inhibiting YAP activator previously identified by our group, synergistically activates TEAD-LUC reporter activity as determined by changes in Bliss independence calculations^36^ (Fig. S1E, F). The observed synergistic effect indicates that sCMF231 may act orthogonally to or in unison with an ANXA2-based mechanism to regulate Hippo signaling.

**Figure 1.**
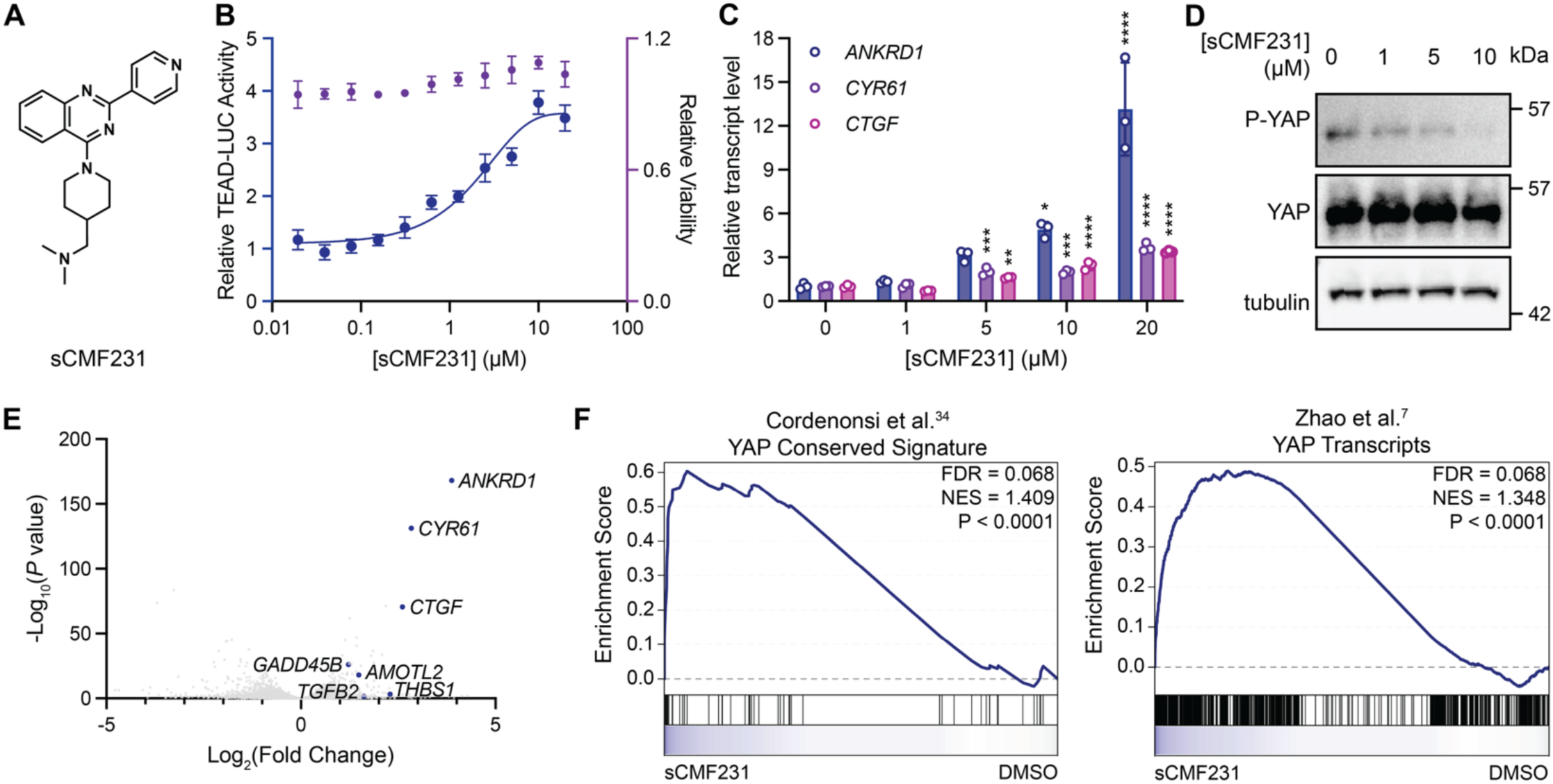
sCMF231, a pharmacological activator of YAP-driven transcription. (A) Structure of sCMF231. (B) Relative luminance values and viability after 24h treatment of 293A-TEAD-LUC cells with the indicated concentrations of sCMF231 (n=3 biological replicates, mean and s.d.). (C) Relative transcript levels of YAP-dependent genes (*ANKRD1, CYR61,* and *CTGF*) from HEK293A cells treated for 24h with sCMF231, measured by RT-qPCR (n=3, mean and s.d.). Statistical significance was determined using one-way ANOVA per transcript (*p<0.05, **p<0.01, ***p< 0.001, ****p<0.0001). (D) Western blot analysis of anti-phospho-YAP (P-YAP) and total YAP levels from HEK293A cells treated with the indicated concentrations of sCMF231 for 6h. (E) Volcano plot representing transcripts identified by RNA-sequencing from 293A cells treated for 24h with sCMF231. Labeled points indicate YAP target transcripts. (F) GSEA plots of YAP-dependent curated gene sets (Cordenonsi et al.^34^ (MSigDB: M2871) and Zhao et al.^7^) from 293A cells treated for 24h with sCMF231 (10 µM).

### FUBI is the relevant cellular target of sCMF231

To identify the relevant intracellular target of sCMF231, we first generated a photo-activatable affinity probe, mCMN927, which bears a photo-crosslinking diazirine moiety as well as a bio-orthogonal alkyne reactivity handle (Fig. 2A). Importantly, mCMN927 retained the capacity to augment TEAD-LUC reporter activity with a similar potency to sCMF231 (EC_50_ = 2.1 µM; Fig. 2B). To survey the potential targets that are covalently labeled in response to probe treatment, HEK293A cells were treated with mCMN927 for 1 hour, crosslinked with 365 nm light, and subjected to click-reaction-based conjugation of rhodamine-azide, revealing two predominant bands around 30 kDa that were selectively labeled at 1 µM (Fig. S3A). Liquid chromatography-tandem mass spectrometry (LC-MS/MS) analysis of excised bands following click-reaction-based conjugation of biotin-azide and enrichment with streptavidin pulldown revealed the protein of interest to be the precursor fusion protein Finkel-Biskis Reilly murine sarcoma virus-associated ubiquitously expressed (FAU). Using purified recombinant FAU protein, we found that mCMN927 could dose-dependently label FAU protein *in vitro* with an EC_50_ of 4.3 µM (Fig. S3B, C). To further confirm FAU as the relevant target of sCMF231, we knocked down FAU in HEK293A cells by approximately 50% using two different lentiviral shRNAs, both of which increased levels of YAP-controlled transcripts *ANKRD1*, *CYR61,* and *CTGF* to a similar magnitude as treatment with sCMF231 (Fig. 2C, D). Conversely, transient overexpression of FAU in HEK293A cells decreased the YAP-dependent transcriptional response (Fig. S4A). Upon 48-hour shRNA knockdown of FAU, changes to the transcriptome by the two shRNAs strongly mimicked that of sCMF231 treatment as indicated by DESeq analysis and GSEA of this RNA-seq data (Fig. 2E, S4B). Strikingly, these shRNAs also robustly enacted a gene set curated from the RNA-seq profile of sCMF231 by GSEA (Fig. 2F), strongly supporting the notion that FAU knockdown can recapitulate the transcriptional effect of sCMF231 treatment.

**Figure 2.**
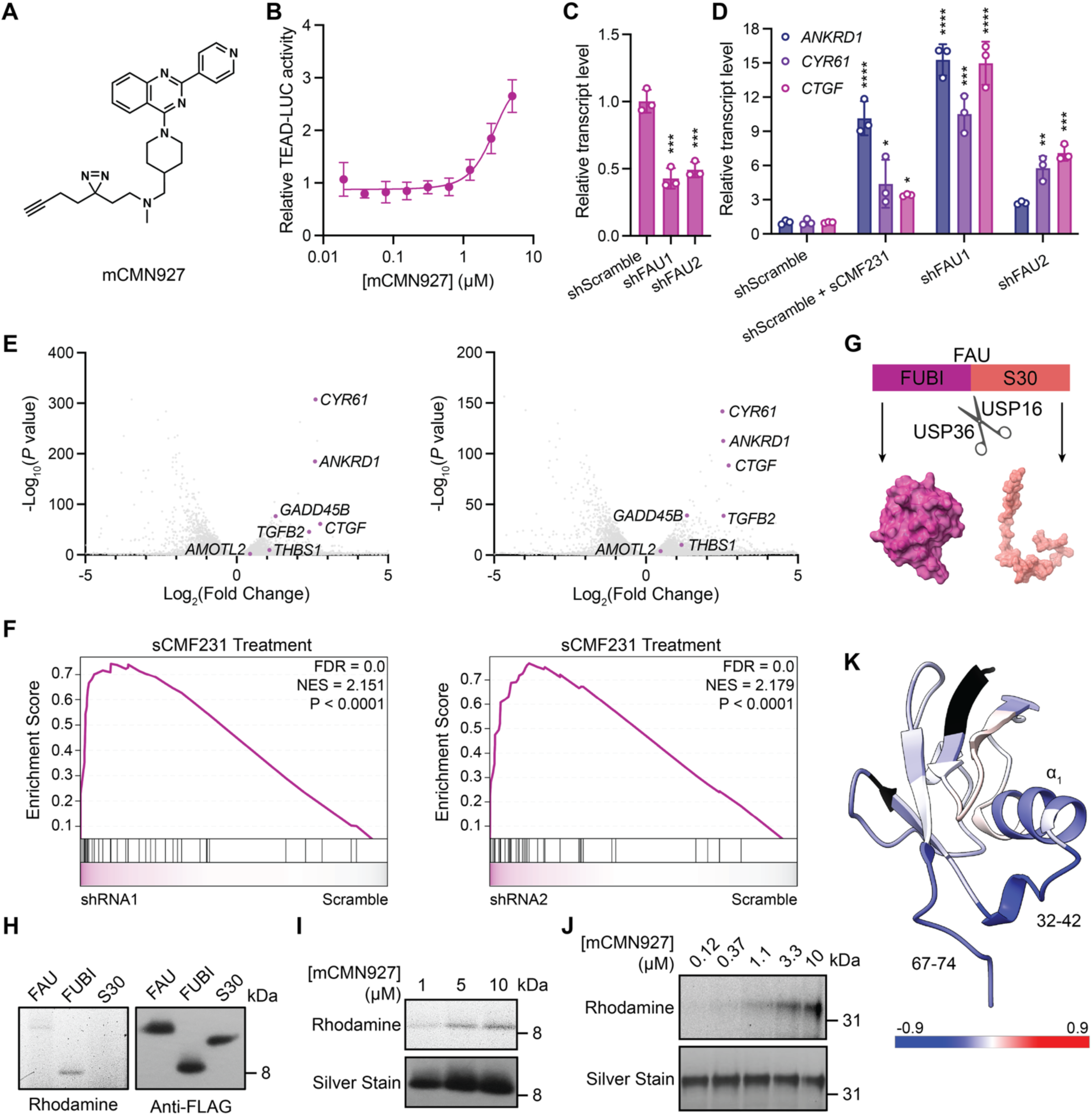
sCMF231 targets FUBI to activate YAP. (A) Structure of photo-activatable affinity probe mCMN927. (B) Relative luminance values after 24h treatment of 293A-TEAD-LUC cells with the indicated concentrations of mCMN927 (n=3 biological replicates, mean and s.d.). (C) Relative transcript levels of FAU following knockdown by the indicated shRNA, measured by RT-qPCR (n=3, mean and s.d.). (D) Relative transcript levels of YAP-dependent genes (*ANKRD1, CTGF, CYR61*) expressing the indicated shRNA, or treated with 10 µM sCMF231 for 24h, measured by RT-qPCR (n=3, mean and s.d.). (E) Volcano plot representing transcripts identified by RNA-sequencing from 293A cells expressing FAU shRNA1 (left) or FAU shRNA2 (right). Labeled points indicate YAP target transcripts. (F) GSEA plot of a sCMF231-treated curated gene set from 293A cells expressing FAU shRNA1 (left) or FAU shRNA2 (right). (G) Schematic of FAU cleavage by USP16 and USP36 to yield the ubiquitin-like protein FUBI (PDB: 2L7R) and ribosomal protein S30 (PDB: 4UG0). (H) Representative fluorescent gel scan and anti-FLAG Western blot analysis of rhodamine-azide labeled material after FLAG-IP of FLAG-FAU, FLAG-FUBI, and FLAG-S30. (I) Representative fluorescent gel scan and anti-FLAG Western blot analysis of rhodamine-azide labeled material after FLAG-IP of FLAG-FUBI with 1h treatment of 1, 5, or 10 µM mCMN927. (J) Representative fluorescent gel scan of rhodamine azide-based labeling of recombinant GST-FUBI exposed to mCMN927. (K) Solution NMR-structure of FUBI (PDB:2L7R) displaying the fractional uptake differences of deuterium from HDX-MS between untreated FUBI and FUBI treated with 600 µM sCMF231. A darker blue color indicates less deuterium uptake in the treated versus untreated sample. Black indicates amino acids with no coverage. Statistical analyses in (C, D) are one-way ANOVAs (*p<0.05, **p<0.01, ***p<0.001, ****p<0.0001).

FAU is a fusion precursor protein that is post-translationally cleaved by the ubiquitin proteases USP16 and USP36 to yield the ubiquitin-like protein FUBI (also known as MNSFβ) and ribosomal protein S30^37–39^ (Fig. 2G). We next sought to understand which of these two protein products, FUBI or S30, was the relevant target of sCMF231. We transiently overexpressed N-terminally FLAG-tagged constructs of FAU (FLAG-FAU), FUBI (FLAG-FUBI), or S30 (FLAG-S30) in HEK293T cells for 48 hours and then treated with 10 µM of mCMN927 followed by click-reaction-based conjugation of rhodamine-azide. Labeling experiments revealed that mCMN927 labeled the FLAG-FAU and FLAG-FUBI transgenes but not FLAG-S30 (Fig. 2H). mCMN927 also labeled FLAG-FUBI *in situ* in a dose-dependent manner (Fig. 2I). In addition, labeling of FLAG-FUBI by mCMN927 was competed by a molecular excess of free sCMF231 (Fig. S3D). Furthermore, recombinant preparations of FUBI, but not S30, were dose-dependently labeled by mCMN927 with an EC_50_ of 3.5 µM (Fig. 2J, S3E). Lastly, to investigate the nature of the interaction of sCMF231 with FUBI, we performed a hydrogen/deuterium exchange mass spectrometry (HDX-MS) experiment^40^. sCMF231 treatment reduced deuterium exchange within two key regions: residues 32-42 (EGIAPEDQVVL), comprising the linker between the central α-helix and the β-sheet, and residues 67-74 (VAGRMLGG), which comprises the C-terminal tail of FUBI, including the diglycine motif required for covalent conjugation (Fig. 2K, S5A-E). These data suggest that sCMF231 likely binds FUBI at or near residues 32-42, perhaps creating a conformational change to the protein that allosterically alters the positioning of the C-terminal tail.

### Fubylation of ANXA2 regulates Hippo Pathway activity

FUBI is an understudied ubiquitin-like protein that contains a C-terminal diglycine motif like ubiquitin, putatively endowing the protein with the potential to be conjugated to proteins through isopeptide linkages^41^. Indeed, some studies have suggested that FUBI, like ubiquitin, can be covalently conjugated to signaling proteins such as Formyltetrahydrofolate Dehydrogenase, Bcl-G, and Endophilin II^42–44^. Since FUBI has no annotated targets that are known Hippo pathway regulators, we sought to identify putative protein targets that might be conjugated to FUBI using affinity purification mass spectrometry. We employed a two-step affinity purification protocol using the tandem histidine-biotin (HB) tag, which contains (starting from the N-terminus) a 6-HIS sequence, Biotin carboxyl carrier protein (BCCP) for *in situ* biotinylation, as well as a TEV cleavage site flanked by flexible linker regions, ending with FUBI. This tag allows for sequential purification by Ni^2+^ and streptavidin resins^45^ under fully denaturing conditions to specifically preserve only covalent FUBI interactors (Fig. 3A). Accordingly, we generated an FAU transgene with an N-terminal HB tag, which upon cleavage in cells would yield free HB-FUBI capable of precipitation by sequential purification. An HB-FUBI transgene itself could also be expressed, but HB-FAU consistently expressed at higher levels (not shown). We transiently overexpressed HB-FUBI in HEK293T cells, followed by tandem purification with Ni-NTA resin and streptavidin agarose, observing several bands detectable by silver staining and anti-Streptavidin Western blotting that were robustly enriched (Fig. S6A). Gel bands excised from purified material identified many potential targets of FUBI, including Annexin A1 (ANXA1) and ANXA2, S100-A proteins (obligate ANXA protein family binders), and desmosome components Desmoglein 1, Desmoplakin, and Plakoglobin. Among these, ANXA2 immediately stood out as a potential target of relevance, as we have previously shown that ANXA2 is a druggable component of the Hippo pathway^15^.

**Figure 3.**
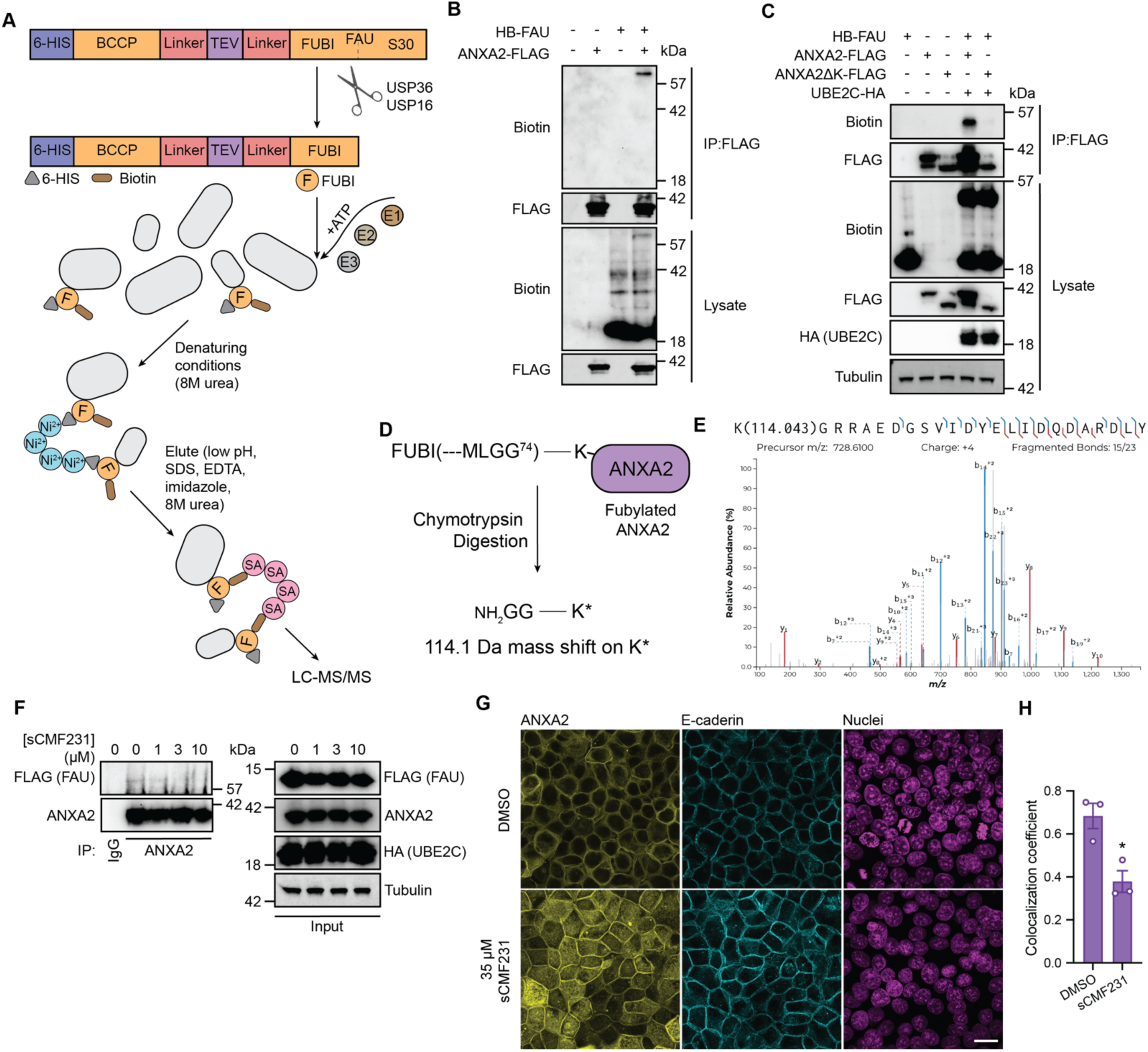
Fubylation of ANXA2 regulates its cellular localization. (A) Schematic of HB-FAU with a N-terminal 6-HIS for Ni-NTA resin purification and BCCP (Biotin Carboxyl Carrier Protein) for *in situ* biotinylation. HB-FAU gets cleaved *in situ* to yield HB-FUBI. Through the activity of an E1 activator, an E2 conjugator, and an E3 ligase, HB-FUBI is covalently conjugated to target substrates. Cell lysates are prepared under fully denaturing conditions and stepwise purified from Ni-NTA resin (Ni^2+^) and streptavidin agarose (SA). Purified samples are analyzed by LC-MS/MS. (B) Immunoblotting analysis of HB-FAU from anti-FLAG immunoprecipitated content from HEK293T cells. (C) Immunoblotting analysis of HB-FAU from anti-FLAG immunoprecipitated content from HEK293T cells transiently overexpressing either wild-type ANXA2 or ANXA2 with no lysines. (D) Schematic of FUBI-conjugated ANXA2 following chymotrypsin digestion containing a diglycine remnant of FUBI covalently attached to a lysine. Amino acids are represented by single-letter code. (E) Sequence and fragmentation pattern (MS/MS spectrum) of the modified peptide produced by chymotrypsin proteolysis of a sample collected from **C**. ANXA2 K176 has the intact diglycine modification. (F) Immunoblotting analysis of FLAG-FAU from anti-ANXA2 immunoprecipitated content from HEK293T cells transiently overexpressing FLAG-FAU and UBE2C-HA and treated with the indicated concentrations of sCMF231 for 24h. (G) Representative images of anti-ANXA2 (yellow) and anti-E-cadherin (teal) immunofluorescent staining (with Hoechst 33342 in pink to visualize nuclei) of MCF7 cells treated with control or 35 µM sCMF231 for 24h (scale bar = 20 µm). (H) Quantification of anti-ANXA2 and anti-E-cadherin correlative immunofluorescent staining. Statistical significance was determined by a univariate two-sided t-test (*p<0.02).

To understand if ANXA2 is indeed an interactor of FUBI, we performed an orthogonal experimental technique involving co-immunoprecipitation studies with FLAG-tagged ANXA2 (ANXA2-FLAG) and HB-FAU transgenes at high cell density (Hippo activating conditions) in HEK293T cells. We found that ANXA2-FLAG precipitated an anti-Biotin and anti-FLAG positive ANXA2-FUBI adduct at ∼60 kDa, a weight corresponding roughly to the mass of ANXA2 (∼37 kDa) and FUBI (∼20 kDa with the HB tag, Fig. 3B). We additionally performed a similar experiment in which ANXA2-FLAG and HB-FAU were co-expressed in HEK293T cells and then purified using streptavidin enrichment in fully denaturing conditions. Notably a similar ∼55 kDa anti-biotin and anti-FLAG positive mass was observed in denaturing conditions, consistent with the notion that the interaction between FUBI and ANXA2 is covalent in nature (Fig. S6B). To determine if fubylation, akin to ubiquitination, occurs on a lysine residue of ANXA2, we generated an ANXA2-FLAG transgene in which every lysine is mutated to arginine (ANXA2ΔK-FLAG). Anti-FLAG immunoprecipitation experiments determined that ANXA2ΔK-FLAG transgene was unable to form the ∼55 kDa adduct like ANXA2-FLAG transgene, indicating ANXA2 is fubylated on a lysine residue (Fig. 3C). To confirm the covalent nature of this modification, we performed post-translational modification analysis via mass spectrometry (PTM-MS). Ubiquitin’s C-terminus of amino acid residues LRLRGG contrasts with FUBI’s C-terminus of GRMLGG. Therefore, we employed cleavage by chymotrypsin, which results in a -GG remnant on the target protein only via fubylation and not by ubiquitination (Fig. 3D). This cleavage contrasts with the commonly used trypsin digestion, which results in -GG remnants on ubiquitinated proteins, but would create an -MLGG remnant for FUBI. We performed PTM-MS on an excised band ∼55 kDa from an enrichment experiment in which ANXA2-FLAG and HB-FAU were co-expressed in HEK293T cells and then purified using streptavidin enrichment in fully denaturing conditions. Indeed, this band contained both ANXA2 and FUBI tryptic fragments (with 40 ANXA2 peptides and ∼40% sequence coverage of ANXA2), and a -GG chymotryptic remnant was identified on K176 of ANXA2 by MS/MS (Fig. 3E). Finally, we sought to understand if sCMF231 treatment alters the interaction between ANXA2 and FUBI. The precipitation of transiently overexpressed FLAG-FAU from HEK293T cells by endogenous ANXA2 was inhibited by 24-hour treatment with sCMF231 (Fig. 3F). Collectively, these results confirm that ANXA2 can be fubylated in cells and that treatment with sCMF231 decreases the efficiency of this fubylation event in some way.

Poly-ubiquitination is often a signal to promote proteasomal degradation of modified proteins. However, FUBI lacks most internal lysine residues that are present in ubiquitin, including those that canonically serve as poly-ubiquitination substrate hubs, ubiquitin K48 and K63 (Fig. S7). In fact, the only lysine in FUBI is K25 (corresponding to ubiquitin K27), which is not solvent accessible in the structures of ubiquitin or FUBI, and no structures of K27-linked polyubiquitin chains exist^46^. Unlike polyubiquitination, which is associated with proteasomal degradation, mono-ubiquitination typically regulates the localization and cellular activity of proteins^47^. Previous work on ANXA2 suggests that ANXA2 regulates Hippo pathway activity by binding to YAP and shepherding it to the plasma membrane to receive inhibitory phosphorylation, with delocalization of ANXA2 away from the plasma membrane serving as a strong Hippo inactivating signal. We therefore evaluated whether sCMF231 alters the intracellular localization of ANXA2. Consistent with previous work, we found that ANXA2 localizes to the plasma membrane of MCF7 cells in at high cell density (Fig. 3G). However, in the presence of sCMF231 (35 µM, 24 hours), ANXA2 was found to delocalize from the plasma membrane, demonstrating an almost two-fold loss of colocalization with membrane protein E-cadherin (Fig. 3G, H). We also evaluated endogenous ANXA2 levels in HEK293A cells following 24-hour treatment with sCMF231 and found that up to 50 µM treatment of compound does not affect ANXA2 levels, indicating that fubylation most likely does not regulate ANXA2 stability (Fig. S6C). These data indicate that fubylation of ANXA2 regulates the localization of ANXA2 to the plasma membrane and that inhibition of FUBI conjugation could likely drive ANXA2 membrane delocalization.

### UBA1, UBE2C, APC11 are the writers of ANXA2 fubylation

Conjugation of ubiquitin and ubiquitin-like proteins (Ubls) to their target proteins is dependent on the successive activities of 3 classes of “writer” proteins: activating enzymes (E1), conjugating enzymes (E2), and ligating enzymes (E3)^48^. As there is no previous evidence that FUBI is conjugated to target proteins, these conjugation enzymes for FUBI have not yet been elucidated^49^. To fully understand the mechanism of how FUBI regulates ANXA2 and how this regulation changes with sCMF231 treatment, we set out to identify the enzymes required for ANXA2 fubylation.

To identify potential fubylation machinery, we again used the tandem affinity purification experiment to look for FUBI interactors. HEK293T cells were plated for 48 hours transiently overexpressing HB-FUBI, and the total sample of tandem affinity-purified HB-FUBI was analyzed using LC-MS/MS. From the 383 proteins identified as putative covalent FUBI interactors, fewer than 10 were known ubiquitination “writing” enzymes (Fig. 4A). We first focused on ubiquitin activating enzyme 1 (UBA1), one of the E1 enzymes for writing ubiquitin modifications^50^. UBA1 activates ubiquitin through the formation of a high-energy thioester linkage between the ubiquitin C-terminus and UBA1’s active site cysteine with the use of ATP. To validate UBA1 as the FUBI activating enzyme, we first assessed if it could be charged by FUBI in cells. We performed co-immunoprecipitation studies with FLAG-tagged UBA1 (UBA1-FLAG) and HB-FAU. UBA1-FLAG precipitated FUBI in a covalent manner, as FUBI migrated to ∼130 kDa, the approximate molecular weight of UBA1-FLAG and FUBI with the HB tag (Fig. 4B). Similarly, HB-FAU was found to precipitate UBA1 via streptavidin enrichment in fully denaturing conditions, again indicating a covalent interaction (Fig. 4C). To parse whether the activation of FUBI by UBA1 is mediated through this thioester bond in a similar manner to ubiquitin, we conducted an *in vitro* charging assay with recombinant FUBI, His-tagged UBA1 (UBA1-HIS), and ATP. We found that in the presence of ATP, UBA1 is charged with FUBI, as indicated by the presence of the high molecular weight band ∼120 kDa in the Anti-FUBI Western blot, the approximate molecular weight of UBA1-HIS and untagged FUBI (Fig. 4D). Importantly, this interaction is competed away through the addition of beta-mercaptoethanol (βME), indicating the interaction between UBA1 and FUBI is indeed mediated through an activating thioester bond.

**Figure 4.**
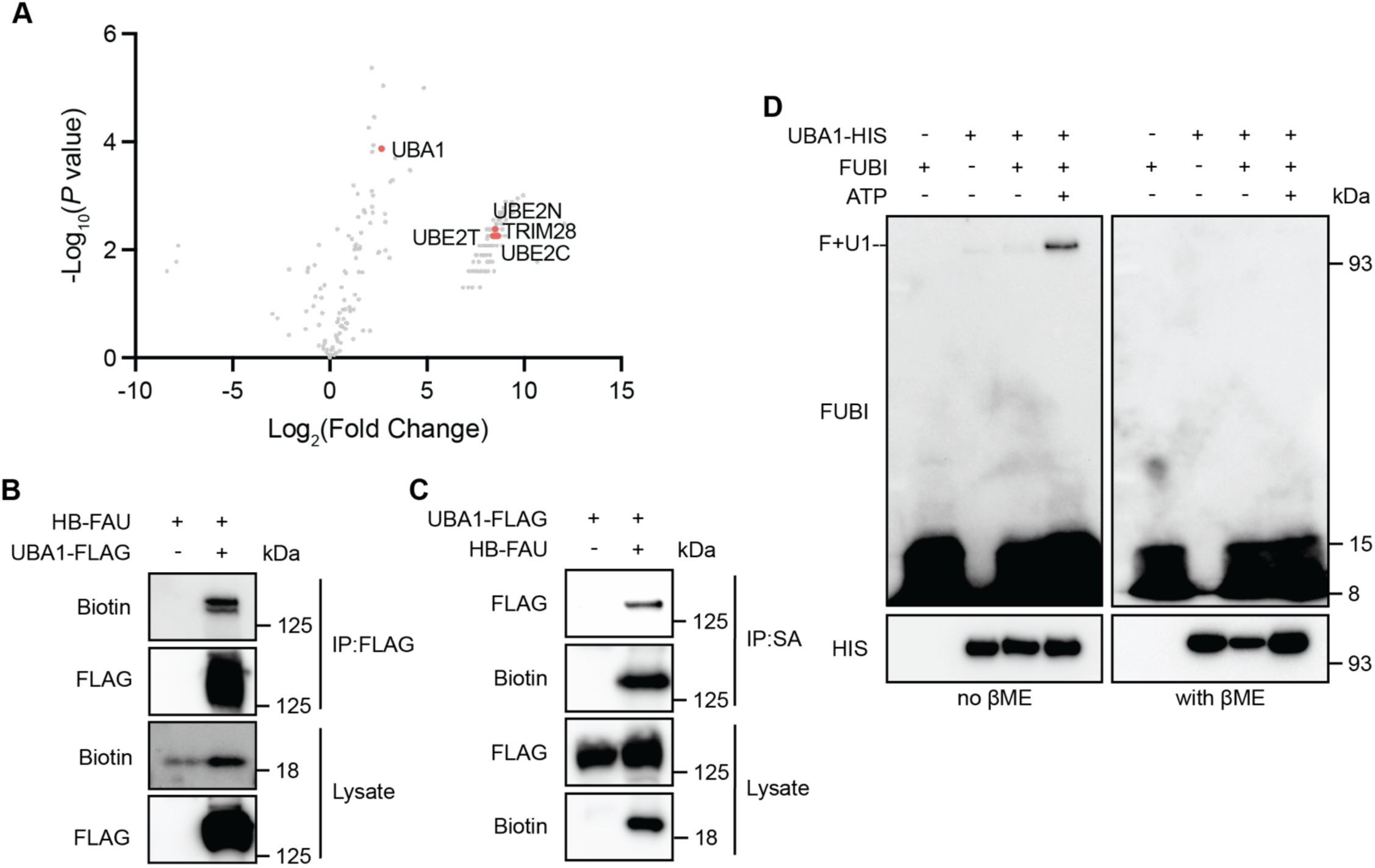
UBA1 is the FUBI activating protein. (A) Volcano plot of proteins enriched by streptavidin from HEK293T cells transiently overexpressing HB-FUBI. Marked points indicate known ubiquitination machinery identified. (B) Immunoblotting analysis of HB-FAU from anti-FLAG immunoprecipitated content from HEK293T cells. (C) Immunoblotting analysis of UBA1-FLAG from anti-streptavidin immunoprecipitated content from HEK293T cells. (D) Anti-FUBI Western blot showing the interaction of UBA1 and FUBI *in vitro* that is competed by βME.

We next focused on identifying the relevant E2 enzyme. Of the three E2 enzymes identified from the tandem affinity purification proteomics experiment (Fig. 4A; UBE2C, UBE2N, and UBE2T), only lentiviral shRNA mediated knockdown of UBE2C was found to increase the levels of YAP-controlled transcripts *ANKRD1, CYR61,* and *CTGF* (Fig. S8A, B). This data indicates that while other E2 enzymes might be putative FUBI writers, only UBE2C regulates Hippo pathway activity. We then confirmed the capacity of UBE2C to covalently load FUBI in cells. From experiments involving the co-expression of FLAG-tagged UBE2C (UBE2C-FLAG) and HB-FAU in HEK293T cells, we found that UBE2C and FUBI interact in a covalent manner, as the interaction of a ∼40 kDa UBE2C-FUBI adduct could be identified both by FLAG immunoprecipitation as well as by streptavidin enrichment in fully denaturing conditions (Fig. 5A, S8C). To determine the importance of UBE2C in fubylating ANXA2, we co-expressed UBE2C-FLAG, ANXA2-FLAG, and HB-FAU and evaluated FUBI conjugation after streptavidin enrichment in denaturing precipitation conditions. The expression of UBE2C-FLAG markedly augmented the FUBI modification of ANXA2 so that it could be strongly precipitated by streptavidin (Fig. 5C). We note that two potential fubylated ANXA2 products are precipitated, one stronger band ∼55 kDa and one weaker band ∼65 kDa. To determine if the interaction between UBE2C and FUBI is mediated through the transthiolation reaction of FUBI from UBA1 to UBE2C’s active site cysteine, we conducted an *in vitro* charging assay with recombinant FUBI, UBA1-HIS, His-tagged UBE2C (UBE2C-HIS), and ATP. *In vitro*, in the presence of ATP, UBA1 transfers FUBI to UBE2C, as indicated by the band ∼28 kDa in the anti-FUBI Western blot (Fig. 5B). Additionally, the interaction between FUBI and UBE2C is competed away through the addition of βME, indicating the transfer of FUBI from UBA1 to UBE2C is mediated through modification of the active site cysteine (Fig. S8D). Since sCMF231 decreases the fubylation of ANXA2 in cells, we next investigated if this decrease might be due to a change in the FUBI-charging capacity of UBE2C. Indeed, *in vitro*, the charging of UBE2C with FUBI significantly decreases in a concentration-dependent manner with sCMF231 treatment (Fig. 5D, S8E). Finally, we confirmed the interaction between UBE2C and ANXA2 with a co-immunoprecipitation experiment involving expression of ANXA2-HA and UBE2C-FLAG in HEK293T cells, as we could identify this interaction via anti-FLAG or anti-HA-based immunoprecipitation by Western blotting (Fig. 5E, S8F). Collectively, these data suggest that UBE2C is necessary and sufficient for fubylation of ANXA2 and that sCMF231 likely inhibits ANXA2 conjugation by decreasing the efficiency of sCMF231 charging.

**Figure 5.**
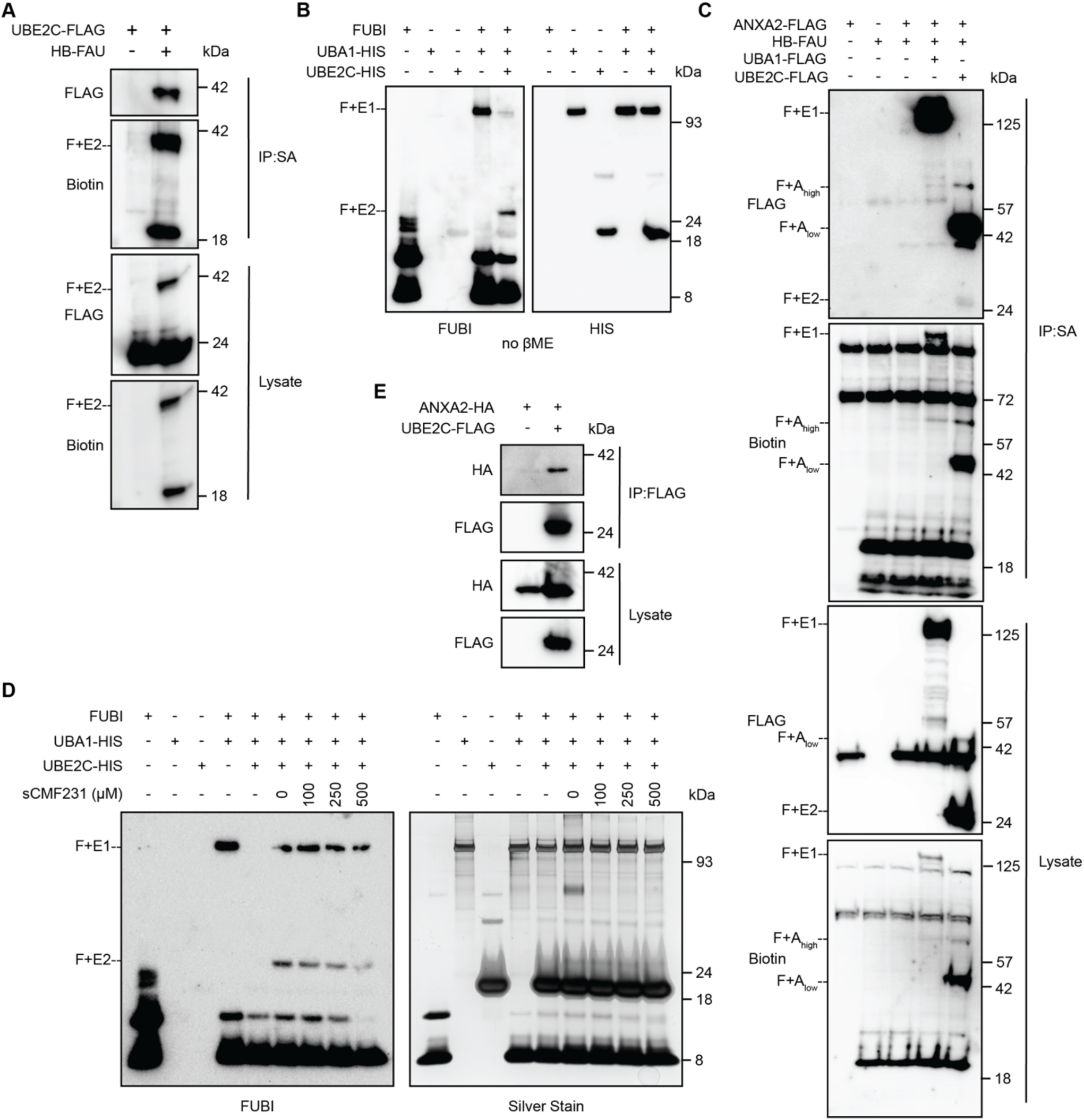
FUBI requires the activity of UBE2C for conjugation to ANXA2. (A) Immunoblotting analysis of UBE2C-FLAG from anti-streptavidin immunoprecipitated content from HEK293T cells. (B) Anti-FUBI Western blot showing the interaction of UBE2C and FUBI *in vitro*. (C) Immunoblotting analysis of UBA1-FLAG, UBE2C-FLAG, and ANXA2-FLAG from anti-streptavidin immunoprecipitated content from HEK293T cells. (D) Anti-FUBI Western blot and silver stain showing that the interaction of UBE2C and FUBI decreases in response to sCMF231 treatment *in vitro*. (E) Immunoblotting analysis of ANXA2-HA from anti-FLAG immunoprecipitated content from HEK293T cells. (A-D) F+E1 indicates the covalent complex of FUBI with UBA1. F+E2 indicates the covalent complex of FUBI with UBE2C. F+A_low_ indicates the stronger (lower molecular weight) fubylated ANXA2 band. F+A_high_ indicates the weaker (higher molecular weight) fubylated ANXA2 band.

We finally turned our attention to identifying the relevant E3 writer of fubylation. E3s generally fall into two classes: 1) homologous to the E6AP carboxyl terminus (HECT) domain family E3s that make a thioester intermediate with the active site cysteine of the E3 and 2) really interesting new gene (RING) and RING-related E3s that mediate the direct transfer of ubiquitin and Ubls from E2 to substrate^51,52^. We identified only one E3 ligase in our tandem affinity purification experiment and we subsequently found it is not the relevant E3 for ANXA2 fubylation (not shown). Since the majority of E3 ligases do not make covalent bonds with ubiquitin, we hypothesized fubylation may be written by a RING E3 ligase. UBE2C canonically acts as the E2 of the anaphase promoting complex/cyclosome (APC/C) which controls the poly-ubiquitination and degradation of various mitotic components to advance the cell cycle. The APC/C complex is a macromolecular machine of 14 subunits^53^ with the catalytic module consisting of APC2, the cullin subunit, and APC11, the RING domain subunit^54^. To understand the potential activity of APC/C in ANXA2 fubylation, we focused on this E3 catalytic module. Knocking down APC2 and APC11 using lentiviral shRNA increased the levels of YAP controlled targets *ANKRD1, CYR61,* and *CTGF* (Fig. 6A). In addition, 24-hour treatment of HEK293A cells with proTAME, an inhibitor of the APC/C complex^55^, moderately induced these YAP-controlled transcripts (Fig. S9). Through a co-immunoprecipitation experiment with APC11-HA, we found that APC11 was able to precipitate ANXA2-FLAG (Fig. 6B). Lastly, to evaluate the importance of APC11 in the fubylation of ANXA2, we overexpressed APC11-HA in the co-immunoprecipitation experiment of HB-FAU by ANXA2-FLAG and found that, while not nearly as strong as UBE2C overexpression due to low expression levels of APC11, APC11 enhanced the precipitation of FUBI (Fig. 6C). This data points to APC/C as a relevant E3 ligase capable of writing fubylation on ANXA2.

**Figure 6.**
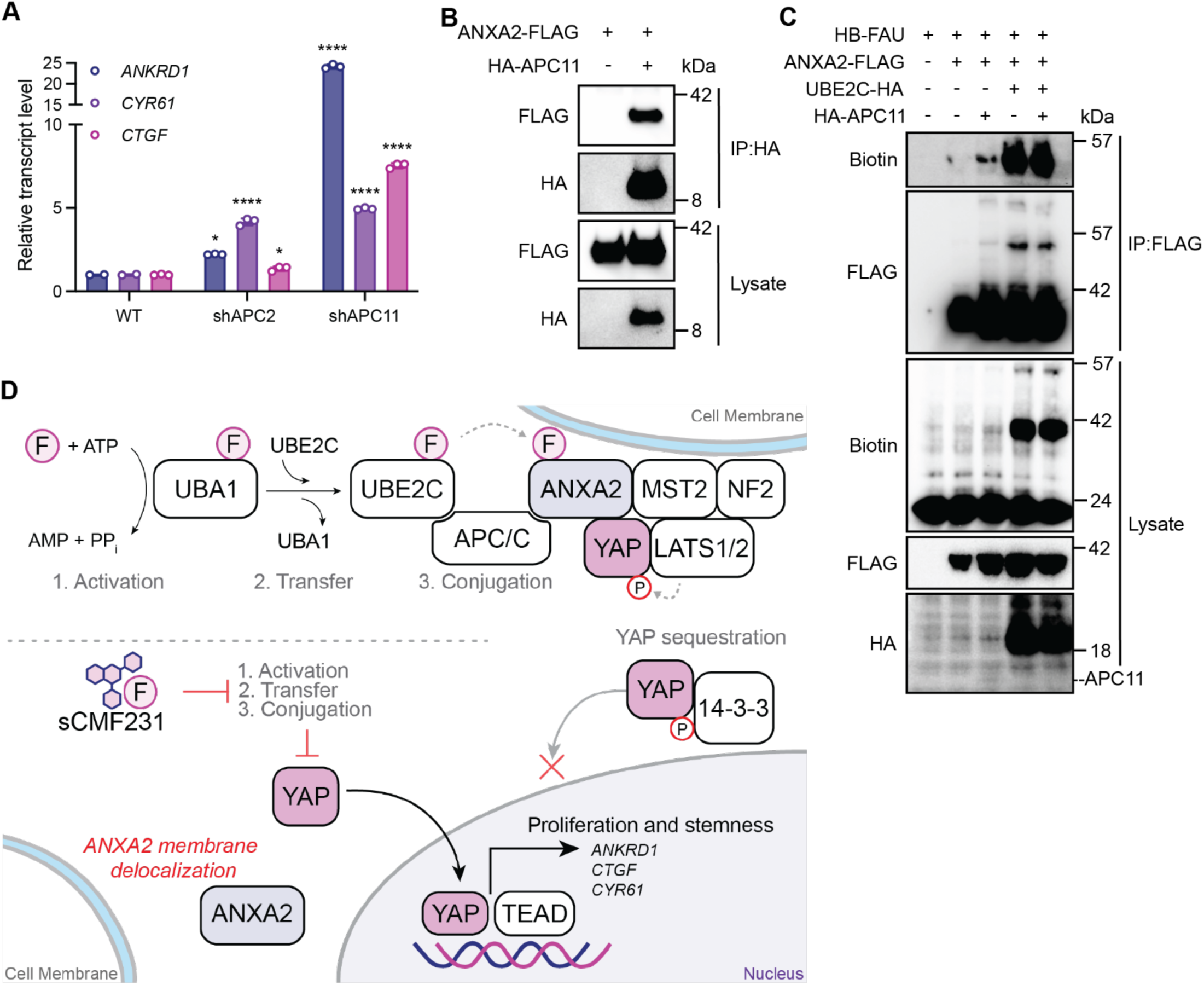
APC/C is an E3 ligase for ANXA2 fubylation. (A) Relative transcript levels of YAP-dependent genes (*ANKRD1, CTGF, CYR61*) expressing the indicated shRNA, measured by RT-qPCR (n=3, mean and s.d.). Statistical significance was determined by one-way ANOVA (*p<0.05, ****p<0.0001). (B) Immunoblotting analysis of ANXA2-FLAG from anti-HA immunoprecipitated content from HEK293T cells. (C) Immunoblotting analysis of ANXA2-FLAG, UBE2C-HA, and HA-APC11 from anti-FLAG immunoprecipitated content from HEK293T cells. (D) A mechanistic model depicting that in the unperturbed system, FUBI is covalently conjugated to ANXA2 via the activation by UBA1, transfer to UBE2C, and conjugation with APC/C. Fubylated ANXA2 localizes at the membranes, bringing YAP in contact with core kinases of the Hippo pathway to receive inhibitory phosphorylation, resulting in its cytoplasmic retention. Upon treatment with sCMF231, FUBI does not activate, transfer, or conjugate as efficiently, leading to a lack of ANXA2 fubylation. Unmodified ANXA2 cannot localize YAP to the membrane, leaving YAP free to enter the nucleus, interact with TEADs, and activate its transcripts.

## Discussion

To uncover additional context specific inhibitors of Hippo pathway signaling, we report here the discovery of sCMF231, a quinazoline containing small molecule that elicits a YAP-activating response in cells without inhibiting the core Hippo pathway kinases MST1/2 and LATS1/2. Notably, among YAP activating small molecules reported to date, sCMF231 enacts the most specific YAP transcriptional program, as we have shown by GSEA. Chemical proteomic target identification experiments revealed the target of the compound to be FUBI, a ubiquitin-like protein that is poorly studied and has historically been thought by a majority of the field to be incapable of covalent conjugation to target proteins. We demonstrate that loss of the FUBI encoding transcript FAU specifically recapitulates the effect of sCMF231, and we further delineate how FUBI conjugation modulates Hippo pathway activity. The discovery of sCMF231 adds another small molecule to the pharmacological toolkit for probing and understanding Hippo pathway regulation. Further work uncovering the spectrum of other cell types in which sCMF231 elicits a YAP-activating response will help determine how broadly FUBI acts as a regulator of Hippo pathway signaling and undoubtedly aid in future efforts to potentially target FUBI for regenerative medicine-based applications.

To our knowledge, sCMF231 is the first compound that directly engages a ubiquitin-like protein. As a small globular protein with no obvious ligand binding pocket, it is perhaps unsurprising that small molecules of this kind have yet to be identified. Instead, most modulators of ubiquitination function by blocking various “writing” processes versus targeting the protein directly^56–59^. A notable exception is BRD1732, a cytotoxic molecule that depletes the supply of ubiquitin monomers available for conjugation by directly alkylating the C-terminus of ubiquitin itself^60^. Unlike BRD1732, sCMF231 was found to non-covalently target FUBI, supported by experimentation *in vitro* and *in situ* with a photo-activatable probe of sCMF231. Compound binding to FUBI was further reinforced by HDX-MS data which revealed two peptides of FUBI that considerably decrease in their deuterium exchange capacity in response to sCMF231 treatment. One of these fragments, residues 67-74, comprises the C-terminal tail of FUBI, which we have shown is capable of conjugation to client proteins. As E2s recognize the C terminus of ubiquitin within the juxtaposition of the E1 and E2 active sites^61–63^, it is conceivable that sCMF231 may conformationally change how FUBI fits into and is recognized at this interface, and thereby decrease efficiency of E2 charging, as we saw with UBE2C. A critical limitation of this work is that we have not fully determined the nature of sCMF231 binding to FUBI. Co-crystal structures as well as additional biophysical work will undoubtedly shed more light on this interaction, potentially helping to understand how sCMF231 specifically engages FUBI relative to other ubiquitin-like proteins as well as provide the basis to develop more potent FUBI ligands.

Our data indicates that FUBI regulates Hippo pathway activity by covalently modifying ANXA2, and loss of this modification promotes the delocalization of ANXA2 from the membrane thereby activating YAP (Fig. 6D). sCMF231 is not the first small molecule that alters the localization of ANXA2 to modulate Hippo pathway activity. We recently identified the small molecule PY-60, a small molecule that directly targets ANXA2, decreasing its capacity for membrane association^15^. In this context, YAP loses capacity for LATS1/2 catalyzed phosphorylation at the plasma membrane, but, by staying bound to ANXA2, is also actively dephosphorylated by the PP2A complex. Several lines of inquiry are required to better understand how fubylation controls YAP activity through ANXA2. Proteomics and co-immunoprecipitation experiments investigating changes in ANXA2’s interactome with and without sCMF231 treatment will yield insights to when, within this signaling node, fubylation of ANXA2 occurs and if the same downstream signaling process, like PP2A association, also occur when ANXA2 delocalizes from the membrane. Further, since the FAU gene is a constitutively expressed gene, it will be important to identify the upstream signals that trigger fubylation of ANXA2 to limit YAP activity. Finally, an examination of changes in ANXA2 ubiquitination upon fubylation could be useful in providing a fuller picture of how post-translational modifications govern ANXA2’s activity. Differences in ANXA2 ubiquitination may indeed account for the presence of ANXA2-FUBI conjugates that appear at distinct molecular weights (∼55 kDa and ∼65 kDa). In all, the identification of fubylation of ANXA2 as a robust and specific regulator of YAP activity indicates ANXA2 is perhaps even more central to the regulation of the Hippo pathway than previously appreciated and supports its importance as a druggable component of this pathway.

FUBI has been proposed as capable of modifying proteins in a limited number of reports, but excitingly, our work is the first to provide unequivocal proof of covalent conjugation of FUBI to a specific lysine on ANXA2 through - GG proteomics. Recent work has focused on deubiquitinases (DUBs) of FUBI, identifying both USP16 and USP36 as DUBs that cleave FAU into mature FUBI and S30 and are potential de-fubylases, or “erasers”, for FUBI conjugates^37,38^. This study adds essential information to this emerging field, as we have identified the first “writers” of ANXA2 fubylation: UBA1, UBE2C, and APC/C. Mechanistically, our data indicates sCMF231 blocks the efficient transfer of FUBI from UBA1 to UBE2C, decreasing fubylation of ANXA2, as indicated through *in vitro* fubylation assays and endogenous co-immunoprecipitation experiments. Unlike other Ubls, which possess their own E1 enzyme, FUBI is unique in that it uses UBA1 as its activating enzyme, one of the two activating enzymes used by ubiquitin^50^. Mechanistic and structural studies will be critical in providing insights into how UBA1 is charged by FUBI and if sCMF231 might additionally interfere with this process. We identified UBE2C as an E2 capable of serving as a conjugating enzyme for FUBI and one that strongly regulates Hippo pathway activity, as indicated by the large increase in YAP-dependent transcripts upon UBE2C knockdown. A proteomics experiment from another study also identified UBE2C as an important regulator of the Hippo pathway, but its mechanistic role was not further studied^64^. While other E2 enzymes interact with FUBI based on our proteomic experiments, UBE2C is the only E2 that both regulates YAP activity and increases the levels of fubylated ANXA2 *in situ*. Whether the other E2s identified in this work serve as a bona fide FUBI writers will be of key importance to future work in uncovering the breadth by which fubylation regulates cellular signaling. As with UBA1, a co-crystal structure of FUBI bound to UBE2C will help provide further mechanistic insight into how recognition of these two proteins occurs as well as how sCMF231 interferes with this process.

As there are over 600 E3 ligases, the relevant E3 ligase for ANXA2 fubylation was more difficult to parse. Since our proteomic studies in fully denaturing conditions did not yield E3 ligase interactors of FUBI that facilitate ANXA2 fubylation, we focused our attention on known, non-covalent RING-type E3 ligases that use UBE2C as the E2. Of the two most studied E3 ligases for UBE2C (APC/C and SAG)^65^, knockdown of APC/C catalytic components APC2 and APC11 surprisingly induced an increase in YAP-controlled transcripts. APC/C is canonically thought of as a regulator of the cell cycle, acting with co-factors CDH1 and CDC20 to trigger and maintain different phases of the cell cycle^66^. Previous work has indicated that APC/C positively regulates YAP activity during cell cycle progression by targeting LATS1/2 for degradation^67^. Our data contradict this reported activity, as knockdown of APC/C catalytic components instead activates YAP. Interestingly, there is evidence that APC11 is able to ubiquitinate substrates without the rest of the APC/C subunits^68^. Necessarily, we will need to query whether APC11 itself or the APC/C complex as a whole mediates the fubylation of ANXA2. Some data point towards APC11 acting as the E3 ligase on its own: 1) APC11 knockdown greatly increases the expression of YAP target transcripts, whereas APC2 knockdown is less effective; 2) APC/C inhibitor proTAME, which functions by inhibiting binding of APC/C by adaptors Cdh1 and CDC20, only very mildly upregulates YAP target transcripts. These results provide evidence that perhaps the activity of the complex as a whole is not required for controlling YAP activity. Knockdown studies of APC/C components that are not part of the catalytic component but integral to APC/C activity and substrate recognition, such as APC10, will provide key data to better understand the requirement of the entire APC/C complex^69^. Verification of the interaction between FUBI, ANXA2, and these various APC/C subunits will also be required. Adding sCMF231 to co-immunoprecipitation experiments will further contribute to understanding the importance of these interactions with FUBI. However, reconstitution of the proposed system *in vitro* ultimately will validate which components of APC/C are necessary and sufficient to fubylate ANXA2. Importantly, the cellular location where ANXA2 is fubylated remains undetermined. ANXA2, while mostly cytoplasmic, can be detected in the nucleus in a cell-cycle dependent manner, specifically accumulating during the G1 phase^70^. Active APC/C^Cdh1^ can also be found in the nucleus of G1 cells^71^. Therefore, it is conceivable that ANXA2 is fubylated in the nucleus by APC/C during G1 and is then exported to assert its regulatory activity on the Hippo pathway in the cytoplasm and at the plasma membrane. Cell synchronization experiments via double thymidine block followed by co-immunoprecipitation and co-immunofluorescence will help elucidate when and where ANXA2 is fubylated in the cell and if APC/C plays an essential role. Alternatively, APC11 localizes both to the nucleus and the cytoplasm. Accordingly, it may be possible that fubylation of ANXA2 may occur in the cytoplasm by APC11 without need for the rest of the APC/C^72^. Finally, we must consider that there are other E3 ligases that we did not identify in our proteomics studies or literature search that may be necessary or sufficient for fubylating ANXA2. A CRISPR or siRNA screen against all known E3 ligases to look for changes in YAP activity would narrow the field for potential E3 fubylation ligases.

Much remains to be explored in the potential fubylation of other proteins. Existing literature has cultivated a list of fubylated target proteins, including Formyltetrahydrofolate Dehydrogenase, Bcl-G, and Endophilin II, among others^42–44^. Our proteomics experiments also identified a host of other potential fubylated proteins, a resource that will hopefully help define the cellular fubylome. These proteins included other annexin proteins and their binders, S100-A proteins. Evaluating the potential of other annexin family members to be fubylated could aid in the understanding of potential redundant signaling mechanisms. Furthermore, we identified multiple components of desmosomes as FUBI targets, specialized cell-cell adhesion junctions that link to the cytoskeleton and give mechanical strength to tissues^73^. Desmosomes have also been specifically linked to YAP regulation^74,75^. These data indicate that FUBI may also covalently modify desmosome components to nucleate several different Hippo regulatory nodes at the membrane to control YAP phosphorylation. Currently, only one commercial antibody exists towards FUBI, and in our hands, it does not readily label endogenous fubylated substrates. Ongoing efforts to optimize a potent antibody that targets both monomeric and conjugated forms of FUBI will better help define the spectrum of proteins targets capable of being fubylated.

Here, we have serendipitously identified fubylation as a druggable ubiquitin-like system that controls Hippo pathway activity. In this context, ANXA2 is covalently modified by FUBI, a ubiquitin like protein that co-opts traditional ubiquitin machinery, UBA1, UBE2C, and APC/C. We posit that fubylated ANXA2 more aptly associates with membranes, where it shepherds YAP to receive inhibitory phosphorylation from the MST1/2 and LATS1/2 kinase cascade. sCMF231 inhibits the efficiency of ANXA2 fubylation, resulting in ANXA2 delocalization away from the membrane and subsequent loss of YAP phosphorylation. Identifying fubylation as a central component governing Hippo pathway activity underscores the power of pharmacology and chemical genetics to uncover novel signaling regulators that might not be recognized without specific chemical tools.

## Materials and Methods

### Cell Culture

293A-TEAD-LUC reporter cells (HEK-293A-8xGTII-Luc) were a kind gift from the laboratory of Xu Wu (Mass General). HEK293A cells were from Thermo Fisher. MDCK, MCF7, and HEK293T cells were obtained from the American Type Culture Collection (ATCC). HaCaT immortalized human keratinocytes were from AddexBio. HEK293T UBE2C knockout cells were from Abcam. All cell lines were cultured in DMEM (Corning, 10-013-CV) supplemented with 10% FBS (Gibco, 10438-026) and 1% penicillin/streptomycin (Gibco, 15070063) at 37°C and 5% CO_2_. MCF7 cells were additionally supplemented with 0.01 mg/mL insulin (Sigma, I0516) and 1% non-essential amino acids (Gibco, 11140050).

### Miniaturized Reporter Assays

293A-TEAD-LUC cells (2,500) were plated in 50 µL of growth medium without FBS in white 384-well plates (Corning). 48h later, 100 nL of compound solvated in DMSO was then transferred to each well using a Bravo Automated Liquid Handling Platform (Agilent) outfitted with a pintool head. For activity assays, 30 µL of BrightGlo reagent solution (Promega, diluted 1:3 in water) was added to each well after 24 hours of compound treatment. For cytotoxicity, 30 µL of CellTiter-Glo reagent solution (Promega, diluted 1:6 in water) was added to each well after 72 hours of compound treatment. Luminescence values were recorded on an Envision plate reader (Perkin Elmer). For testing compounds in combination, the Bliss expectation (C) was calculated using the equation *C* = (*A* + *B*) – (*A × B*), where *A* and *B* represent the fractional luminescence of two compounds at a given dose. The difference between the observed luminescence of both compounds in combination and the calculated Bliss expectation is plotted as “Δ Bliss independence.” Values greater than zero for this calculation represent a response which is greater than additive (synergistic).

### shRNA Knockdown Studies

Lentiviruses were generated in HEK293T cells by transient expression of the vectors with psPAX2 and pMD2.G packaging vectors. Viral supernatants were collected after 48 hours of expression and passed through a 45 µm syringe filter before exposure to target cells.

| Vector Name | Transgene | Source |
| --- | --- | --- |
| APC2 shRNA | ANAPC2-targeted shRNA | Sigma-Aldrich, TRCN0000004357 |
| APC11 shRNA | ANAPC11-targeted shRNA | Sigma-Aldrich, TRCN0000038800 |
| FAU shRNA #1 | FAU-targeted shRNA | Sigma-Aldrich, TRCN0000011075 |
| FAU shRNA #2 | FAU-targeted shRNA | Sigma-Aldrich, TRCN0000342246 |
| UBE2C shRNA #1 | UBE2C-targeted shRNA | Sigma-Aldrich, TRCN0000349490 |
| UBE2C shRNA #2 | UBE2C-targeted shRNA | Sigma-Aldrich, TRCN0000368929 |
| UBE2C shRNA #3 | UBE2C-targeted shRNA | Sigma-Aldrich, TRCN0000368994 |
| pLKO1-shYAP2 (YAP shRNA) | YAP-targeted shRNA | Addgene, plasmid #27369 |
| Scramble | Non-targeting shRNA | Sigma-Aldrich, SHC002 |
| pMD2.G | VSVG, lentiviral packaging | Addgene, plasmid #12259 |
| psPAX2 | Gag-Pol, lentiviral packaging | Addgene, plasmid #12260 |

### Quantitative Reverse Transcription PCR (qRT-PCR)

After 48 h of transgene expression and/or 24 h of compound treatment, cells were trypsinized and collected by subsequent centrifugation at 1200 x*g*. RNA was isolated using an RNeasy kit (Qiagen) and RNA concentrations were determined using a NanoDrop instrument. 1 µg of RNA was subjected to reverse transcription reaction with High-Capacity cDNA Reverse Transcription Kit (Thermo Fisher). Quantitative RT-PCR reactions were measured on a Viia 7 Real-Time PCR system (Thermo Fisher) using Power SYBR Green (Thermo Fisher, 4367659) and transcript-specific primers below. Reactions were normalized to GAPDH levels for each biological replicate and relative transcript abundance calculated using the comparative C_t_ method.

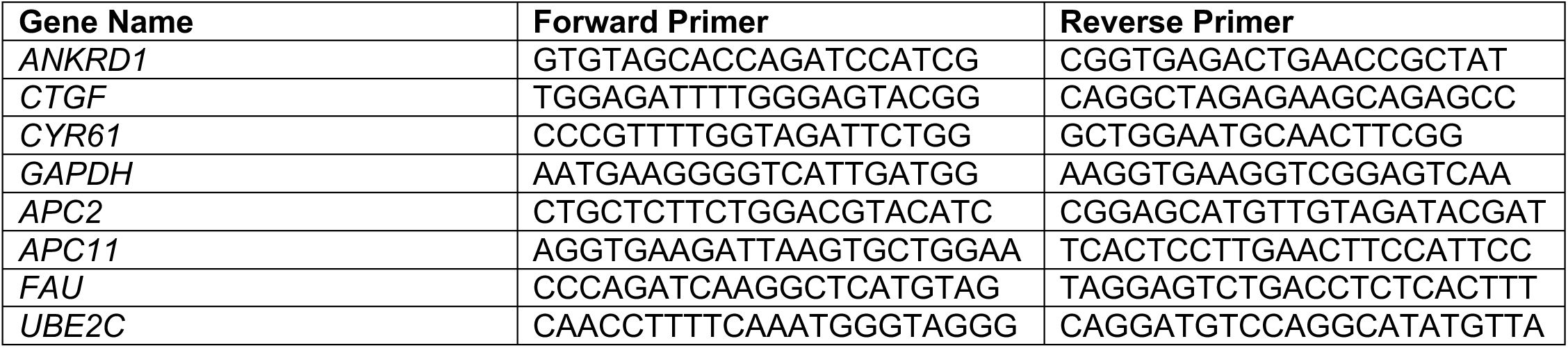

### RNA-Sequencing

For RNA-Seq experiments, cells were treated with the indicated concentration of compound or subjected to shRNA knockdown, and RNA collected as described above. RNA from three biological replicates was sent to BGI for library preparation and sequencing on a DNBSEQ Platform. Paired-end reads were filtered and trimmed with TrimGalore to remove sequencing adapters and low-quality sequences. Reads were then mapped to the human genome (GRCh38) using STAR alignment^76^. Genomic features in mapped reads were counted using featureCounts^77^. Differential gene expression analysis (DESeq2) was carried out using *R*^78^. Gene set enrichment analysis (GSEA) was performed with the desktop Javascript application^79^.

### Cloning

Codon-optimized sequences encoding FLAG-tagged transgenes of N-terminally tagged FAU, FUBI, and S30, and tandem 6-HIS and biotin tags at the N-terminus of FAU and FUBI were obtained from Integrated DNA Technologies as gBlock HiFi Gene Fragments and cloned into the pCMV6 backbone via Gibson assembly using a HiFi DNA Assembly Cloning Kit (NEB). Codon-optimized sequences encoding FAU, FUBI, and S30 were obtained from Integrated DNA Technologies as gBlock HiFi Gene Fragments and cloned into the pGEX-6P-1 GST backbone via Gibson assembly using a HiFi DNA Assembly Cloning Kit (NEB). A codon-optimized sequence encoding 6-HIS-SUMO-FUBI obtained from Integrated DNA Technologies as gBlock HiFi Gene Fragments and cloned into the PET28b backbone via Gibson assembly using a HiFi DNA Assembly Cloning Kit (NEB).

| Vector Name | Transgene | Epitope Tag | Source |
| --- | --- | --- | --- |
| pCMV6-Entry | Empty Vector | None | Origene, PS10001 |
| pGEX-6P-1-GST-FAU | FAU | N-terminal GST | This work |
| pGEX-6P-1-GST-FUBI | FUBI | N-terminal GST | This work |
| pGEX-6P-1-GST-S30 | S30 | N-terminal GST | This work |
| pCMV6-FLAG-FAU | FAU | N-terminal FLAG | This work |
| pCMV6-FLAG-FUBI | FUBI | N-terminal 3x-FLAG | This work |
| pCMV6-FLAG-S30 | S30 | N-terminal 3x-FLAG | This work |
| pCMV6-HB-FAU | FAU | N-terminal 6-HIS-biotin | This work |
| pCMV6-HB-FUBI | FUBI | N-terminal 6-HIS-biotin | This work |
| pCMV6-ANXA2-FLAG | ANXA2 | C-terminal FLAG | Shalhout et al, 2021 |
| pCMV3-ANXA2-HA | ANXA2 | C-terminal HA | Sino Biological, HG13517-CY |
| pCMV6-ANXA2-no lysine | ANXA2 | C-terminal FLAG | This work |
| pCMV6-UBA1-FLAG | UBA1 | C-terminal FLAG | Origene, RC201446 |
| pCMV6-UBE2C-FLAG | UBE2C | C-terminal FLAG | Origene, RC208741 |
| pCMV3-UBE2C-HA | UBE2C | C-terminal HA | Sino Biological, HG16911-CY |
| pET-UbcH10-wt | UBE2C | C-terminal 6-HIS | Addgene, plasmid #8503 |
| 6HIS-SUMO-FUBI | FUBI | N-terminal 6-HIS-SUMO | This work |
| pcDNA3-HA-APC11 | APC11 | N-terminal HA | Addgene, #19900 |

### Immunoblotting

HEK293A cells were plated at 500,000 cells per well in six-well plates. 48 h later, cells were then treated with the indicated concentrations of sCMF231 for 24 h. Cells were washed with PBS and collected with 250 µL of RIPA buffer (EMD Millipore) with scraping. Lysates were clarified by centrifugation (18,253 x*g*, 5 min, 4°C) and protein concentrations determined by absorbance on a NanoDrop instrument (Thermo Fisher). Equal amounts of lysate per sample were mixed with 4x loading dye (2% SDS, 200 mM Tris, 20% glycerol, and 0.01% bromophenol blue) with β-mercaptoethanol (βME) added to a final concentration of 10%. Typically, 30 µg of protein material was resolved by SDS-polyacrylamide gel electrophoresis (SDS-PAGE) in 12-well gels (Invitrogen, 4-12% Bis-tris BOLT gels) in 0.9x MOPS-SDS buffer (Invitrogen). Samples were transferred to polyvinylidene difluoride (PVDF) membrane (Thermo Fisher) using a semi-dry transfer apparatus (Bio-Rad Laboratories). Membranes were blocked at room temperature for 1 h with 5% non-fat dry milk (Bio-Rad Laboratories) in Tris-buffered saline (TBS, Corning) containing 0.1% Tween-20. Primary antibodies were incubated overnight with shaking at 4°C in 5% milk in TBST or 5% bovine serum albumin (BSA) in TBST. Membranes were exposed to either fluorophore-conjugated secondary antibodies (Li-Cor; 1:2000) or HRP-conjugated secondary antibodies (Sigma; 1:3000) in TBST with 5% milk for 30 min (Li-Cor) or 1 h (HRP) at room temperature. Signals were recorded with either a Li-Cor fluorescence imager or exposed to HRP substrate (Dura West Substrate, Pierce) and signals recorded using autoradiography film (Genesee Scientific) or using a ChemiDoc instrument (Bio-Rad).

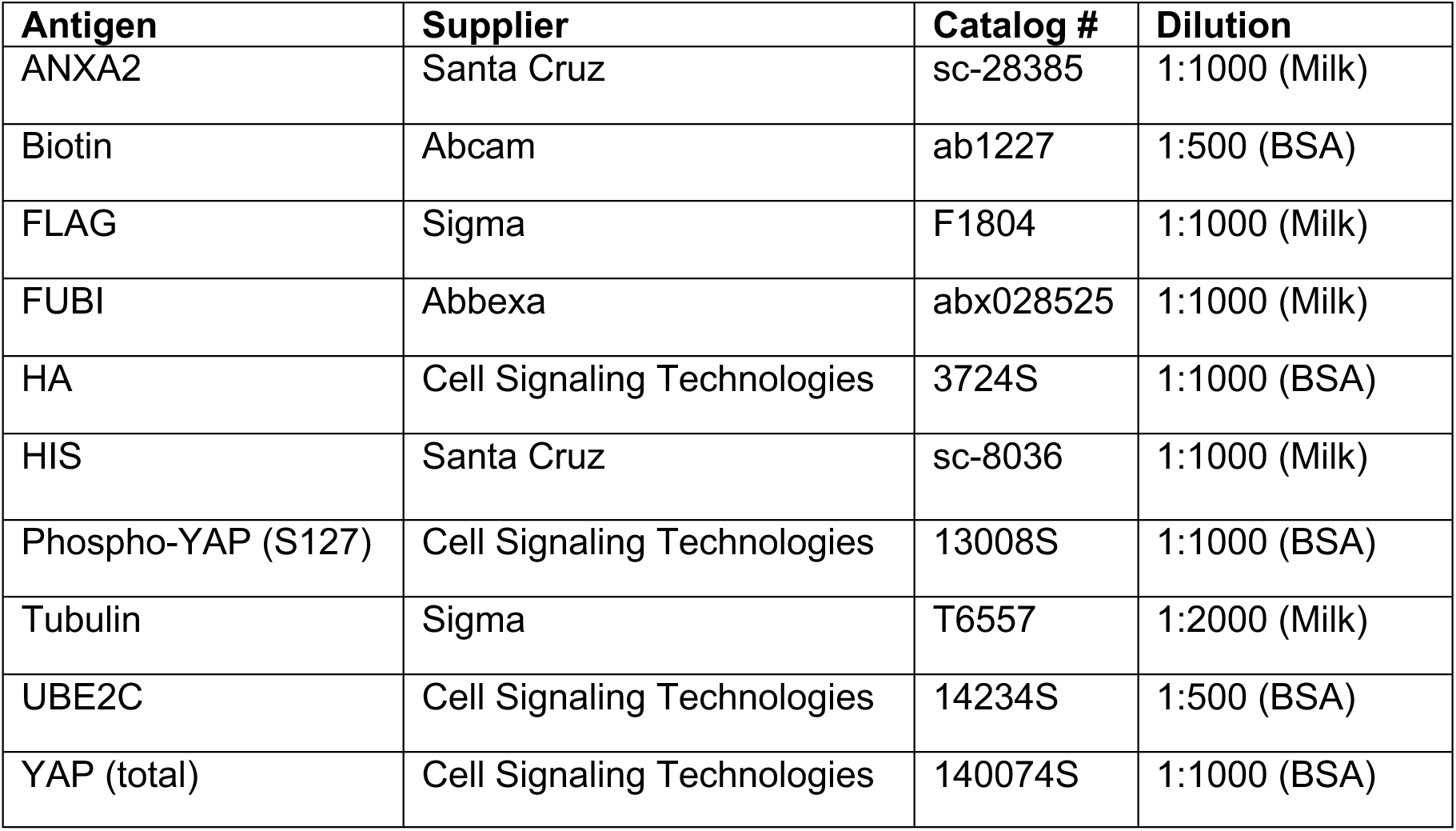

### Target Identification Studies with mCMN927

10 cm tissue culture dishes containing confluent monolayers of 293A cells in standard growth medium were washed twice with PBS and exposed to RPMI medium without phenol red or FBS containing mCMN927 (and sCMF231 for competition experiments) at the indicated concentrations for 1 h at 37°C. Culture dish lids were then removed and the cells exposed to UV-B light (365 nm) for 15 min at 4°C. Cells were washed twice with PBS and scraped into 500 µL of ice-cold PBS. Collected cells were tip sonicated for 5 s and clarified by centrifugation (18,253*g*, 7 min, 4°C). Protein abundance was then measured by absorbance on a NanoDrop instrument (Thermo Fisher). 2 mg of lysate in 1 mL of PBS was exposed to 55 µL of click reaction master mix; 60 µL of a representative mix contains: (1) 30 µL of 1.7 Tris((1-benzyl-1H-1,2,3-triazol-4-yl)methyl)amine (Sigma or Cayman Chemical) in 4:1 tBuOH:DMSO solution, (2) 10 µl of 50 mM CuSO_4_ (Sigma) in water, (3) 10 µl of 5 mM Biotin-PEG3-Azide (Sigma) or 5 mM rhodamine-azide (Sigma no. 760757) in DMSO and (4) 10 µl of 50 mM tris(2-carboxyethyl)phosphine (TCEP) in water. Reactions were vortexed to mix and incubated for 1 h at room temperature in the dark. 500 µL of ice-cold methanol was then added to precipitate the protein content. For imaging, samples were separated in SDS-PAGE Bis-tris gels (Life technologies) and fluorescent signal was recorded with a ChemiDoc imager (Bio-Rad). Gels were Coomassie stained to confirm equal loading. For streptavidin enrichment, protein pellets were resuspended in 1 mL of 0.6% SDS in PBS and added to 200 µL of streptavidin agarose suspended in 10 mL of PBS. After 24 h of rotation at room temperature, beads were washed twice with 0.1% SDS in PBS, twice with PBS, and twice with water. Streptavidin-bound material was eluted by boiling the agarose in 200 µL of 4x SDS sample buffer for 15 min. Immunoblotting for biotinylated protein content was performed as described above, and parallel SDS-PAGE gels were silver stained with a Pierce Silver Stain Kit (no. 24612). For labeling of recombinant protein, protein was diluted to 2 µg per 100 µL of PBS and treated with the indicated concentrations of mCMN927 for 30 min at 37°C. Following exposure to UV-B light for 15 min at 4°C, 10 µL of click reaction master mix was added and incubated for 1 h in the dark at room temperature. Protein was precipitated and washed with 150 µL of ice-cold methanol and samples resuspended in 100 µL of 4x SDS sample buffer. Samples run on SDS-PAGE gels were imaged with a ChemiDoc instrument and silver stained to ensure even loading.

### Immunoprecipitation Studies

For immunoprecipitation studies with epitope-tagged transgenes, plasmids were transiently transfected to HEK293T cells for transgene expression using FuGENE (4 µL FuGENE per 1 µg of DNA) in 100 µL of Opti-MEM (Gibco) per well of a six-well plate (2 µg DNA total per well) or 600 µL of Opti-MEM per 10 cm dish (6-10 µg DNA total per plate). After 48 hours, cells were washed with PBS. For FLAG or HA immunoprecipitation, cells were collected by the addition of 250 µL of RIPA buffer (EMD Millipore) with scraping. For streptavidin immunoprecipitation, cells were collected in fully denaturing lysis buffer (8 M urea, 300mM NaCl, 50mM Tris-HCl, pH = 8, 50mM NaH2PO4, 0.5% NP40) with scraping. Cell lysates were then tip sonicated. Insoluble material was separated by centrifugation (18,253*g*, 7 min, 4°C) and the protein concentration in cellular lysates was determined by absorbance on a nanodrop instrument (Thermo Fisher).

For IP: 1 mg of lysate in 1 mL of lysis buffer was incubated overnight at 4°C with 20 µL of anti-FLAG M2 magnetic bead slurry (MilliporeSigma, no. M8823), overnight at 4°C with 20 µL of anti-HA magnetic bead slurry (Thermo Scientific Pierce, no. 88837), or incubated at room temperature overnight with 50 µL of streptavidin agarose bead slurry (Thermo Scientific Pierce, no. 20353). For FLAG or HA immunoprecipitation, beads were washed three times with lysis buffer, and immunoprecipitated material was eluted with 250 µg per mL of FLAG peptide (DYKDDDDK, Sino Biological) or glycine, pH=2, respectively. For streptavidin immunoprecipitation, beads were washed sequentially with buffer 1 (8 M Urea, 200 mM NaCl, 2% SDS, 100mM Tris, pH 8.0), buffer 2 (8M Urea, 1.2M NaCl, 0.2% SDS, 100mM Tris, 10% EtOH, 10% Isopropanol, pH 8.0), and buffer 3 (8 M urea, 100 NH4HCO3, pH 8). Immunoprecipitated material was then eluted through the addition of 0.1% SDS with boiling. For immunoprecipitation of endogenous proteins, 5 µg of the relevant antibody was incubated with 3 mg of lysate in 1 mL of RIPA overnight at 4°C. 20 µL of Protein G magnetic beads (MedChemExpress no. HY-K0204) were then added to the antibody/lysate solution and incubated for 4h at 4°C. Beads were washed five times with lysis buffer, and immunoprecipitated content was eluted by boiling in 4x sample buffer. Eluted material from immunoprecipitations and whole-cell lysates were evaluated by immunoblotting analyses as described above.

| Antigen | Supplier | Catalog # |
| --- | --- | --- |
| IgG Isotype Control (Rabbit) | Cell Signaling Technologies | 3900S |
| ANXA2 | Proteintech | 11256-1-AP |

### Tandem Affinity Purification Mass Spectrometry

Plasmids were transiently transfected to HEK293T cells for transgene expression using FuGENE (4 µL FuGENE per 1 µg of DNA) in 600 µL of Opti-MEM per 10 cm dish (6 µg DNA total per plate). After 48 hours, cells were washed with PBS and collected in fully denaturing lysis buffer (8 M urea, 300mM NaCl, 50mM Tris-HCl, pH = 8, 50mM NaH2PO4, 0.5% NP40) with scraping. 10 mM imidazole final concentration was added to lysates followed by incubation overnight with Ni-NTA resin (Qiagen, no. 30210). The next day, beads were washed with fully denaturing lysis buffer (pH adjusted to 6.3). Proteins were eluted with buffer B (8M Urea, 200mM NaCl, 50mM Na2HPO4, 2% SDS, 10mM EDTA, 100 mM Tris, 250mM imidazole, pH 4.3) and eluates adjusted back to pH=8. Eluates were then incubated overnight at room temperature with 50 µL streptavidin agarose bead slurry (Thermo Scientific Pierce, no. 20353). The next day, beads were washed sequentially with buffer 1 (8 M Urea, 200 mM NaCl, 2% SDS, 100mM Tris, pH 8.0), buffer 2 (8M Urea, 1.2M NaCl, 0.2% SDS, 100mM Tris, 10% EtOH, 10% Isopropanol, pH 8.0), and buffer 3 (8 M urea, 100 NH4HCO3, pH 8). Washed beads containing affinity-purified proteins were sent to the Sanford Burnham Prebys proteomics core.

Beads were resuspended with 8M urea, 50 mM ammonium bicarbonate. Protein disulfide bonds were reduced with 5 mM tris(2-carboxyethyl)phosphine (TCEP) at 30°C for 60 min, and cysteines were subsequently alkylated with 15 mM iodoacetamide (IAA) in the dark at room temperature for 30 min. Urea was then diluted to 1 M urea using 50 mM ABC, and proteins were subjected to overnight digestion with mass spec grade Trypsin/Lys-C mix (Promega). Following overnight digestion, beads were pulled down and the solution with peptides transferred to a new tube. Finally, samples were acidified with formic acid (FA) and subsequently desalted using AssayMap C18 cartridges mounted on an Agilent AssayMap BRAVO liquid handling system. Cartridges were first conditioned with 100% acetonitrile (ACN) followed by 0.1% FA, samples were then loaded, washed with 0.1% FA, and peptides eluted with 60% ACN, 0.1% FA. The samples were dried down and resuspensed in 2%ACN + 0.1 FA. Peptide quantification was performed using a NanoDrop spectrophotometer (Thermo Scientific).

Samples were then analyzed by LC-MS/MS using a Proxeon EASY-nanoLC system (ThermoFisher) coupled to a Q-Exactive Plus mass spectrometer (Thermo Fisher Scientific). Peptides were separated using an analytical C18 Aurora column (75µm x 250 mm, 1.6 µm particles; IonOpticks) at a flow rate of 200 nL/min using a 120-min gradient: 1% to 5% B in 1 min, 6% to 23% B in 72 min, 23% to 34% B in 45 min, and 34% to 48% B in 2 min (A= FA 0.1%; B=80% ACN: 0.1% FA). The mass spectrometer was operated in positive data-dependent acquisition mode. MS1 spectra were measured in the Orbitrap in a mass-to-charge (m/z) of 350 – 1700 with a resolution of 70,000 at m/z 400. Automatic gain control target was set to 1 x 10^6 with a maximum injection time of 100 ms. Up to 12 MS2 spectra per duty cycle were triggered, fragmented by HCD, and acquired with a resolution of 17,500 and an AGC target of 5 x 10^4, an isolation window of 1.6 m/z and a normalized collision energy of 25. The dynamic exclusion was set to 20 seconds with a 10 ppm mass tolerance around the precursor.

All mass spectra from were analyzed with MaxQuant software version 1.6.11.0. MS/MS spectra were searched against the Homo sapiens Uniprot protein sequence database (downloaded in Apr 2022) and GPM cRAP sequences (commonly known protein contaminants). Precursor mass tolerance was set to 20ppm and 4.5ppm for the first search where initial mass recalibration was completed and for the main search, respectively. Product ions were searched with a mass tolerance 0.5 Da. The maximum precursor ion charge state used for searching was 7. Carbamidomethylation of cysteine was searched as a fixed modification, while oxidation of methionine and acetylation of protein N-terminal were searched as variable modifications. Enzyme was set to trypsin in a specific mode and a maximum of two missed cleavages was allowed for searching. The target-decoy-based false discovery rate (FDR) filter for spectrum and protein identification was set to 1%.

### Post-Translational Modification Mass Spectrometry

For identification of fubylation on ANXA2 lysines, 2.5 µg DNA Tandem-FAU, ANXA2-FLAG, and UBE2C-FLAG each were transiently transfected to HEK293T cells for transgene expression using 30 µL of FuGENE in 100 µL of Opti-MEM (Gibco) per 10 cm dish (7.5 µg DNA total per plate). After 48 h of incubation, cells were collected in 1 mL of fully denaturing lysis buffer. Cell lysates were then tip sonicated and insoluble material was separated by centrifugation (18,253*g*, 7 min, 4°C). The protein concentration in cellular lysates was determined by absorbance on a nanodrop instrument (Thermo Fisher). 12 mg of lysate in 1 mL of lysis buffer was incubated at room temperature overnight with 200 µL of streptavidin agarose bead slurry (Thermo Scientific Pierce, no. 20353). Beads were then washed sequentially with buffer 1 (8 M Urea, 200 mM NaCl, 2% SDS, 100mM Tris, pH 8.0), buffer 2 (8M Urea, 1.2M NaCl, 0.2% SDS, 100mM Tris, 10% EtOH, 10% Isopropanol, pH 8.0), and buffer 3 (8 M urea, 100 NH4HCO3, pH 8). Immunoprecipitated material was then eluted through the addition of 0.1% SDS with boiling. Samples were concentrated using 3 kDa centrifugal filter units (MilliporeSigma, no. UFC500324). After 4x SDS sample buffer was added, immunoblotting for biotinylated and FLAG-tagged protein content was performed as described above, and parallel SDS-PAGE gels were Coomassie stained. Excised bands were sent to the Scripps Research Multi-omics Core.

The gel bands were de-stained, reduced, and alkylated, and then digested overnight with chymotrypsin at room temperature. The peptides were extracted from the gel bands, dried down, and then resuspended and analyzed on a Fusion Orbitrap tribrid mass spectrometer (Thermo). The digest was injected directly onto a 25 cm, 100 µm ID column packed with BEH 1.7µm C18 resin (Waters). Samples were separated using a Themo Neo Vanquish LC system. Peptide identification was done using IP2 and ProLuCID considering no enzyme specificity and 114.042927 modification at lysine (-GG tail).

### Imaging Studies

For ANXA2 localization studies, MCF7 cells were plated in 12-well plates with glass coverslips at 50,000 cells per well in growth medium and propagated for 7 days cells followed by treatment for 24 hours with 35 µM sCMF231. Growth medium was removed, and cells were fixed with ice-cold methanol for 10 min at -20°C. After washing 5 times with PBS, cells were permeabilized with 0.1% Triton-X in PBS for 20 min at room temperature. Primary antibodies in 5% BSA in PBS were incubated overnight at 4°C. After three washes with PBS, Secondary AlexaFluor-conjugated antibodies (Life Technologies; 1:500 in 5% BSA in PBS) with Hoechst 33342 dye were incubated for 1 h at room temperature in the dark. The wells were washed three more times with PBS and then coverslips were removed and mounted onto glass slides using the ProLong Glass Antifade Mountant (Thermo Fisher). Slides were imaged using a laser-scanning Zeiss confocal LSM 720 microscope and colocalization coefficients calculated using Coloc2 (ImageJ).

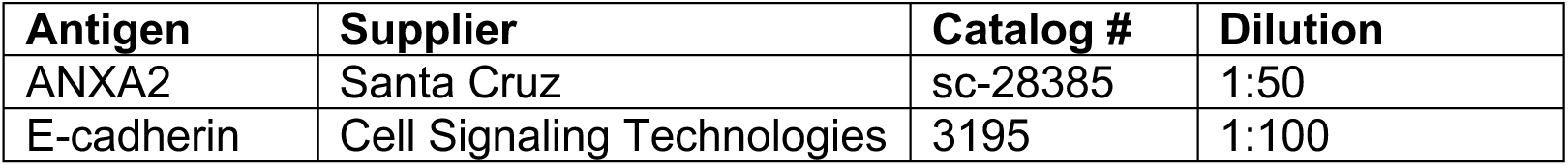

| Antigen | Supplier | Catalog # | Dilution |
| --- | --- | --- | --- |
| ANXA2 | Santa Cruz | sc-28385 | 1:50 |
| E-cadherin | Cell Signaling Technologies | 3195 | 1:100 |

### Human Keratinocyte Proliferation Assays

Immortalized human keratinocytes (HaCaT cells) were plated at 5 x 10^3^ cells per well in 12-well plates in standard growth medium. After 24 h, cells were washed 2 times with PBS, exchanged to reduced-serum-containing medium (DMEM containing 1% penicillin/streptomycin and 0.5% or 2% FBS), and treated with the indicated concentrations of sCMF231. After 7 days of growth under these conditions, cells were fixed in 4% PFA in PBS for 10 min, exposed to 2 µg per mL Hoechst 33342 for 10 min and imaged on a Nikon Eclipse Ti microscope. Nuclei were quantified by an automated ImageJ macro and reported as nuclei per imaging field. Cells were then stained with Rhodamine B for imaging.

### Protein Expression and Purification

For GST-tagged protein expression, BL21 E. coli with the relevant vector were cultured in LB supplemented with Carbenicillin at 37°C until O.D._600_ reached ∼0.6. Expression was induced by adding IPTG to a final concentration of 0.2 mM and cultured for 21 hours at 20°C. Cells were harvested by centrifugation followed by resuspension and lysis by sonication in equilibration buffer (50 mM Tris-HCl, pH=8, 150 mM NaCl). Lysates were clarified by centrifugation and batch bound for 2 hours at 4°C with glutathione agarose (Thermo Scientific Pierce, no. 16100). The solution was transferred to a column, washed with equilibration buffer, and eluted with elution buffer (50 mM Tris-HCl, pH=8, 150 mM NaCl, 10 mM reduced glutathione). Glutathione was removed and TCEP was added to the protein solution using a PD-10 desalting column (Cytiva, no. 17085101). Protein purity was assessed by SDS-PAGE, and purified protein was flash frozen in liquid nitrogen and stored at -80°C.

For FUBI protein expression, an overnight cell culture of BL21 E. coli was grown at 37°C in the presence of Kanamycin. After transfer of the cell culture to 1 L of LB supplemented with Kanamycin, the cell suspension was allowed to reach an O.D._600_ of ∼0.7 before addition of 0.1 mM IPTG (final concentration). Cells were then grown for 21 hours at 18°C and collected by centrifugation. Harvested cells were resuspended in lysis buffer (50 mM sodium phosphate, 300 nM NaCl, 10 mM imidazole, pH=8) and lysed by sonication. Lysates were clarified by centrifugation, and proteins purified by metal ion affinity chromatography by batch binding for 2 hours at 4°C with Ni-NTA resin (Qiagen, no. 30210). The solution was transferred to a column and washed with lysis buffer. 6-HIS-SUMO-FUBI was eluted with an imidazole gradient of elution buffer (50 mM sodium phosphate, 500 nM NaCl, 20-250 mM imidazole). 6-HIS-SUMO-FUBI was then buffer exchanged to cleavage buffer (20 mM Tris-HCl, pH=8, 150 mM NaCl, 1.5 mM DTT). 6-HIS-SUMO was removed from FUBI through the addition of SUMO-Protease/ULP1 (Invitrogen no. 12588-018) at a ratio of 1 mg 6-HIS-SUMO-FUBI : 20 units ULP1. This reaction was allowed to proceed at 4°C overnight. This protein solution was added to Ni-NTA resin for batch binding for 2 hours at 4°C. Protein solution was moved to a column and washed with cleavage buffer. Untagged FUBI was found in the flow through and wash fractions, and protein cleavage and purity were assessed by SDS-PAGE. The protein was further purified by ion exchange chromatography on a Mono Q 5/50 GL column (Cytiva, no. 17-5166-01) with start buffer (20 mM Tris-HCl, pH=8, 1 mM DTT) and elution buffer (20 mM Tris-HCl, pH=8, 1 M NaCl, 1 mM DTT) on a linear gradient from 50-500 mM NaCl. Protein purity was analyzed by SDS-PAGE and intact mass spectrometry, and protein concentration determined by reducing agent compatible BCA (Thermo Scientific Pierce no. 23252). Purified FUBI was flash frozen in liquid nitrogen and stored at -80°C.

For ANXA2 protein expression, BL21 E. coli were cultured in TB supplemented with Carbenicillin at 37°C until O.D._600_ reached ∼0.7-0.9. Expression was induced by adding IPTG to a final concentration of 1 mM and cultured for 14 hours at 30°C. Cells were harvested by centrifugation followed by resuspension and lysis by sonication in equilibration buffer (20 mM Tris-HCl, pH=8, 150 mM NaCl, 10 mM imidazole, 0.1 mM TCEP). Lysates were clarified by centrifugation and batch bound for 2 hours at 4°C with Ni-NTA resin. The solution was transferred to a column, washed with equilibration buffer, and eluted with elution buffer (20 mM Tris-HCl, pH=8, 150 mM NaCl, 250 mM imidazole, 0.1 mM TCEP). Protein was further purified by size exclusion chromatography on a Sephadex 200 increase 10/300 GL column (Cytiva, no. 28990944) with SEC buffer (20 mM Tris-HCl, pH=8, 150 mM NaCl, 2 mM TCEP). After assessment of purity by SDS-PAGE, purified protein was flash frozen in liquid nitrogen and stored at -80°C.

For UBE2C purification, BL21 E. coli were cultured in LB supplemented with Kanamycin at 37°C until O.D._600_ reached ∼0.7. Expression was induced by adding IPTG to a final concentration of 1 mM and cultured for 20 hours at 14°C. Cells were harvested by centrifugation followed by resuspension and lysis by sonication in equilibration buffer (20 mM Tris-HCl, pH=7.4, 136 mM NaCl, 10 mM imidazole, 1 mM DTT). Lysates were clarified by centrifugation and batch bound for 2 hours at 4°C with Ni-NTA resin. The solution was transferred to a column, washed with equilibration buffer, and eluted with imidazole gradient of elution buffer (20 mM Tris-HCl, pH=8, 150 mM NaCl, 20-250 mM imidazole, 0.1 mM TCEP). Protein was further purified by ion exchange chromatography on a Mono Q 5/50 GL column (Cytiva, no. 17-5166-01) with start buffer (20 mM Tris-HCl, pH=7.5, 25 mM NaCl, 0.2 mM TCEP) and elution buffer (20 mM Tris-HCl, pH=7.5, 500 mM NaCl, 0.2 mM TCEP) on a linear gradient from 25-500 mM NaCl. After assessment of purity by SDS-PAGE, purified protein was flash frozen in liquid nitrogen and stored at -80°C.

### In Vitro Fubylation Assays

For UBA1 charging, His-UBA1 (100 nM) (Acro Biosystems, no. UB1-H5248) and FUBI (100 µM) were added to fubylation buffer 1 (final buffer concentration: 20 mM Tris-HCl, pH=8, 5 mM MgCl_2_, 2 mM ATP) to a final volume of 40 µL. The reactions were incubated at 37°C for 1 h and stopped by boiling in SDS sample buffer with and without 10% βME. For UBE2C charging, His-UBA1 (280 nM), UBE2C-HIS (11.5 µM), and FUBI (9.3 µM) were added to fubylation buffer 2 (final buffer concentration: 50 mM HEPES, pH=7.5; 100 mM KCl; 2.5 mM MgCl_2_; 2 mM ATP) to a final volume of 40 µL and incubated for 45 min at 37°C. Reactions were stopped by boiling in SDS sample buffer with and without 10% βME. For fubylation assays with treatment, recombinant proteins were incubated for 30 min at 37°C with the indicated concentrations of sCMF231 before the addition of ATP. Fubylated proteins were visualized by immunoblotting with Anti-FUBI antibody, and an Anti-HIS blot or silver stained gel was used as a protein loading control.

### Hydrogen-Deuterium Exchange Mass Spectrometry (HDX-MS)

HDX-MS was performed at the Biomolecular and Proteomics Mass Spectrometry Facility (BPMSF) of the University California San Diego, using a Waters Synapt G2Si system with HDX technology (Waters Corporation) according to methods previously described^80^. Briefly, deuterium exchange reactions were performed using a Leap HDX PAL autosampler (Leap Technologies). D_2_O buffer was prepared by lyophilizing sample buffer (20 mM Tris-HCl, 100 mM NaCl, 1 mM DTT, pH 8.0) initially dissolved in ultrapure water and redissolving the powder in the same volume of 99.96% D_2_O (Cambridge Isotope Laboratories, Inc.) immediately before use. FUBI (5 μM final) was preincubated for 15 min at RT with 600 μM sCMF231 before storage at 0.1°C before pipetting of exchange reactions. Deuterium exchange was measured in triplicate at each time point (0 min, 0.25min, 0.5 min, 1 min, 2 min, 5 min). For each deuteration time point, 4 μL of protein was held at 25 °C for 5 min before being mixed with 56 μL of D_2_O buffer containing either 3% DMSO (control) or 600 μM sCMF231 (treated). The deuterium exchange was quenched for 1 min at 0.1°C by combining 50 μL of the deuteration reaction with 50 μL of 3M guanidine hydrochloride, final pH 2.66. The quenched sample (90 μL) was then injected in a 100 μL sample loop, followed by digestion on an in-line pepsin column (Immobilized Pepsin, Pierce) at 15°C. The resulting peptides were captured on a BEH C18 Vanguard precolumn, separated by analytical chromatography (Acquity UPLC BEH C18, 1.7 µm 1.0 × 50 mm, Waters Corporation) using a 7–85% acetonitrile gradient in 0.1% formic acid over 7.5 min, and electrosprayed into the Waters Synapt G2Si quadrupole time-of-flight mass spectrometer. The mass spectrometer was set to collect data in the Mobility, ESI+ mode; mass acquisition range of 200–2000 (m/z); scan time 0.4 s. Continuous lock mass correction was accomplished with infusion of leu-enkephalin (m/z = 556.277) every 30 s (mass accuracy of 1 ppm for calibration standard).

For peptide identification, the mass spectrometer was set to collect data in mobility-enhanced data-independent acquisition (MS^E^), mobility ESI+ mode instead. Peptide masses were identified from triplicate analyses and data were analyzed using the ProteinLynx global server (PLGS) version 3.0 (Waters Corporation). Peptide masses were identified using a minimum number of 250 ion counts for low energy peptides and 50 ion counts for their fragment ions; the peptides also had to be larger than 1,500 Da. The following cutoffs were used to filter peptide sequence matches: minimum products per amino acid of 0.2, minimum score of 7, maximum MH+ error of 5 ppm, and a retention time RSD of 5%. In addition, the peptides had to be present in two of the three ID runs collected. The peptides identified in PLGS were then analyzed using DynamX 3.0 data analysis software (Waters Corporation). Peptides containing mutated residues were manually assigned. The relative deuterium uptake for each peptide was calculated by comparing the centroids of the mass envelopes of the deuterated samples with the undeuterated controls following previously published methods^81^. The peptides reported on the coverage maps are actually those from which data were obtained. For all HDX-MS data, at least 2 biological replicates were analyzed each with 3 technical replicates. Data are represented as mean values +/- SEM of 3 technical replicates due to processing software limitations, however the LEAP robot provides highly reproducible data for biological replicates. The deuterium uptake was corrected for back-exchange using a global back exchange correction factor (typically ∼25%) determined from the average percent exchange measured in disordered termini of various proteins^82^. ANOVA analyses and t tests with a p value cutoff of 0.05 implemented in the program, DECA, were used to determine the significance of differences between HDX data points^83^. Deuterium uptake plots were generated in DECA (github.com/komiveslab/DECA), and the data are fitted with an exponential curve for ease of viewing.

### Synthesis of mCMN927

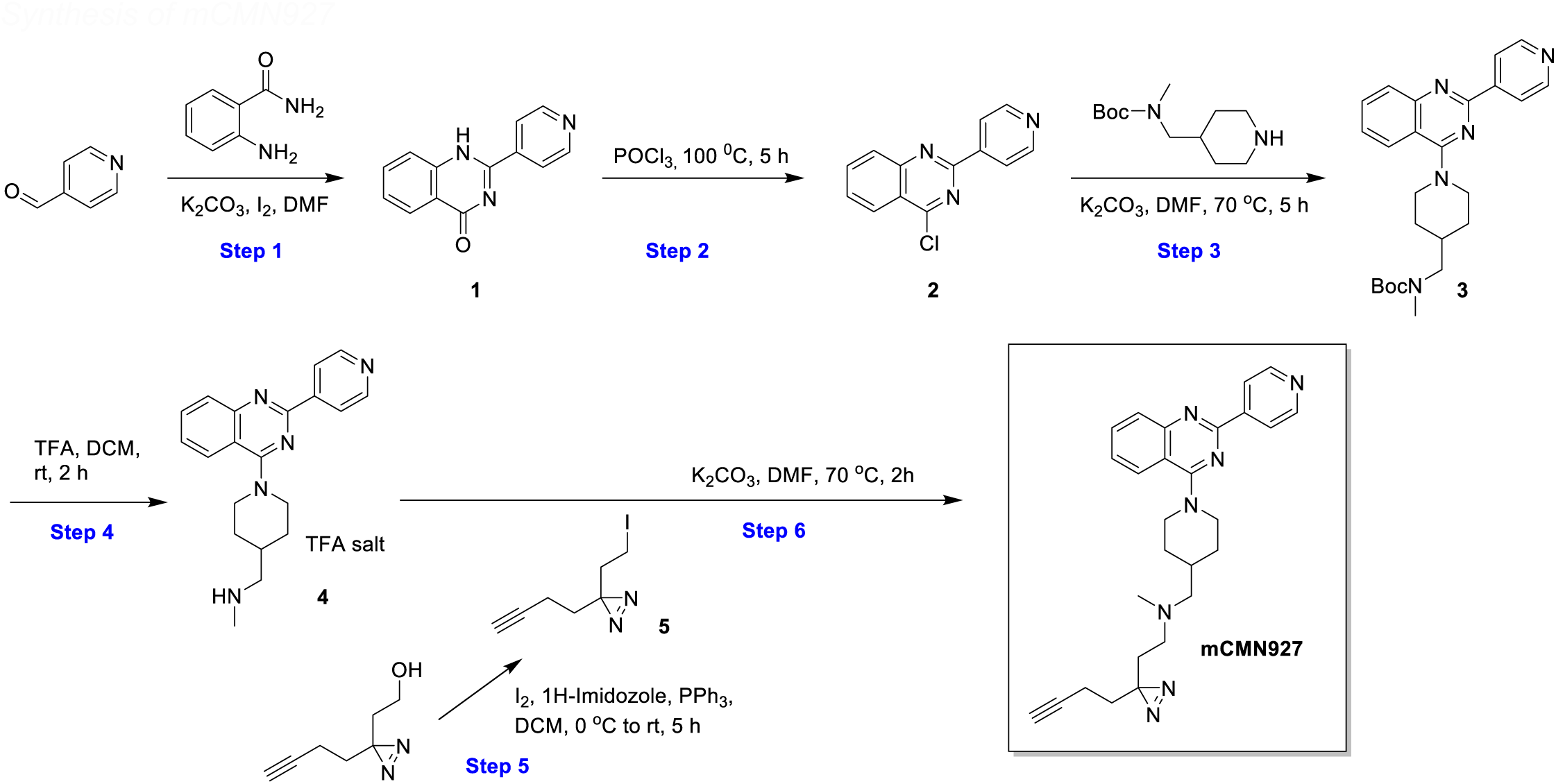

**<u>Step 1</u>**<u>: 2-(pyridin-4-yl)quinazolin-4(1*H*)-one (**1**)</u>: Isonicotinaldehyde (1.0 g, 9.3 mmol, 1.0 eq) was added to a stirred solution of 2-aminobenzamide (1.27 g, 9.3 mmol, 1.0 eq), potassium carbonate (1.28 g, 9.3 mmol, 1.0 eq) and iodine (2.95 g, 11.7 mmol, 1.3 eq) in DMF (20 mL). The reaction mixture was stirred at 80 °C overnight. EtOAc (30 mL) was added, and the layers were separated. The aqueous layer was extracted twice with EtOAc (30 mL) and the combined organic layers were washed with brine (30 mL), dried with Na_2_SO_4_, then the solvent was removed in vacuo to give **1** (1.1 g, 4.9 mmol, yield 52%) as a white solid. MS (ESI): *m/z* calcd. for C_13_H_10_N_3_O^+^ [M+H]^+^ 224.1, found 223.9.

**<u>Step 2</u>**<u>: 4-chloro-2-(pyridin-4-yl)quinazoline (**2**)</u>: A solution of **1** (1.0 g, 4.5 mmol) in POCl_3_ (10 mL) was stirred at 100 °C for 5 hours. The reaction mixture was concentrated and the residue was purified by chromatography on silica gel, eluting (PE:EtOAc = 1:1) to give **2** (700 mg, 2.9 mmol, yield 64%) as a white solid. MS (ESI): *m/z* calcd. for C_13_H_9_ClN_3_^+^ [M+H]^+^ 242.0, found 241.8.

**<u>Step 3</u>**<u>: *tert*-butyl methyl((1-(2-(pyridin-4-yl)quinazolin-4-yl)piperidin-4-yl)methyl)carbamate (**3**)</u>: A solution of **2** (100 mg, 0.41 mmol, 1.0 eq), *tert*-butyl methyl(piperidin-4-ylmethyl)carbamate (177.5 mg, 0.83 mmol, 2.0 eq) and potassium carbonate (114.5 mg, 0.83 mmol, 2.0 eq) in DMF (5 mL) was stirred at 70 °C for 5 hours. EtOAc (15 mL) was added, and the layers were separated. The aqueous layer was extracted twice with EtOAc (15 mL) and the combined organic layers were washed with brine (15 mL), dried with Na_2_SO_4_, then concentrated. The residue was purified by chromatography on silica gel, eluting (PE:EtOAc = 2:1) to give **3** (120 mg, 0.28 mmol, yield 68%) as a yellow solid. MS (ESI): *m/z* calcd. for C_25_H_32_N_5_O_2_^+^ [M+H]^+^ 434.3, found 433.8.

**<u>Step 4</u>**<u>: *N*-methyl-1-(1-(2-(pyridin-4-yl)quinazolin-4-yl)piperidin-4-yl)methanamine (**4**)</u>: To a mixture of **3** (120 mg, 0.28 mmol) in DCM (2 mL) was added TFA (0.4 mL) at 0 °C, and the solution was stirred for 2 hours at room temperature. TLC showed the reaction completed. The reaction mixture was concentrated, and the residue was dissolved in DCM (10 mL). The solution was concentrated again to give crude **4** (110 mg, quantitative) as a white solid (TFA salt) which was used in next step without any further purification.

**<u>Step 5</u>**<u>: 3-(but-3-yn-1-yl)-3-(2-iodoethyl)diazirine (**5**)</u>: To a solution of I_2_ (219 mg, 0.86 mmol, 1.2 eq), 1*H*-imidazole (147 mg, 2.16 mmol, 3.0 eq) and PPh_3_ (207 mg, 2.16 mmol, 3.0 eq) in DCM (5 mL) under nitrogen at 0 °C was added a solution of 2-[3-(but-3-yn-1-yl)diazirin-3-yl]ethanol (100 mg, 0.72 mmol, 1.0 eq) in DCM (1 mL). The reaction mixture was stirred at room temperature for 4 hours. To the reaction mixture was added an aqueous solution of Na_2_S_2_O_3_ (20 mL). The resulting mixture was extracted with EA (3 x 20 mL). The organic phase was washed with saturated brine (20 mL), dried over Na_2_SO_4_, concentrated, and purified by flash column chromatography (PE: EA = 20:1) to obtain **5** (30 mg, 0.12 mmol, yield 17%) as a colorless oil. MS (ESI) – unstable, moved to next step.

**Step 6**: {2-[3-(but-3-yn-1-yl)diazirin-3-yl]ethyl}(methyl)({1-[2-(pyridin-4-yl)quinazolin-4-yl]piperidin-4-yl}methyl)amine (**mCMN927**): A solution of **4** (100 mg, 0.23 mmol, 1.0 eq), diazirine **5** (150 mg, 0.60 mmol, 2.6 eq) and K_2_CO_3_ (124 mg, 0.9 mmol, 3.9 eq) in DMF (3 mL) was stirred under nitrogen at 70 °C for 2 hours. The reaction mixture was filtered, the filtrate was collected and concentrated under pressure. The residue was purified via Genal-Prep-HPLC (Type:X-bridge-C18 150 x 19 mm, 5 µm mobile phase: MeCN-H_2_O (0.05% NH_3_), Ratio: 50-90] to give **mCMN927** (21.5 mg, 0.047 mmol, 21%) as an off-white solid. MS (ESI): *m/z* calcd. for C_27_H_32_N_7_^+^ [M+H]^+^ 454.27, found 453.59; ^1^H NMR (400 MHz, CDCl_3_) δ 8.76 (d, *J* = 5.6 Hz, 2H), 8.39 (dd, *J* = 4.4, 1.6 Hz, 2H), 7.97 (dd, *J* = 8.4 Hz, 0.8 Hz, 1H), 7.91 (d, *J* = 8.4 Hz, 1H) ,7.75-7.73 (m, 1H), 7.49 – 7.44 (m, 1H), 4.52 (d, *J* = 13.2 Hz, 2H), 3.26 – 3.20 (m, 2H), 2.27-2.15 (m, 6H), 2.06 – 1.95 (m, 5H), 1.90-1.82 (s, 1H), 1.68 (t, *J* = 7.6 Hz, 2H), 1.60 (s, 3H), 1.50-1.40 (m, 2H).

### Chemicals

sCMF231 was obtained from Enamine and dissolved in DMSO. proTAME was obtained from Cayman Chemical in methyl acetate. The solvent was evaporated under nitrogen and the compound was re-dissolved in DMSO. No further purification was performed.

### Statistics

EC_50_ values were determined by fitting the fold responses to the mean of DMSO-treated samples using a non-linear regression model in GraphPad Prism. At least three biological replicates were used to determine dose-response curves. Statistical analyses performed in GraphPad Prism were either two-sided t-tests or one-way ANOVA. All experiments were performed at least twice. Data were plotted as means of individually recorded tests and the standard deviation (s.d.) was denoted as error bars where appropriate.

## Acknowledgements

We thank Linh Truc Hoang of the Scripps Research Center for Metabolomics, Jolene Diedrich and Antonio Pinto of the Scripps Research Multi-omics Core, Svetlana Maurya of the Sanford Burnham Prebys Proteomics Core, and Steve Silletti of the University of California at San Diego HDX-MS Core for assistance with mass spectrometry related experiments. We also thank Han Zhou of the Schultz Lab at Scripps Research for advice regarding experimentation for ubiquitin and ubiquitin-like proteins and Thu Nguyen for help optimizing protein expression conditions. This work was supported by the NIH (GM146865 to MJB). MLB was supported by a CIRM training fellowship (EDUC4-12811).

## Author Contributions

M.L.B. and M.J.B. designed research; M.L.B. and E.M.G. performed research; M.L.B., E.M.G., and K.N. analyzed data; N.G.R.D.E. and A.K.C. oversaw the syntheses of small molecules. M.J.B. supervised the project; M.L.B. and M.J.B. wrote the manuscript.

---

**Figure S1.**
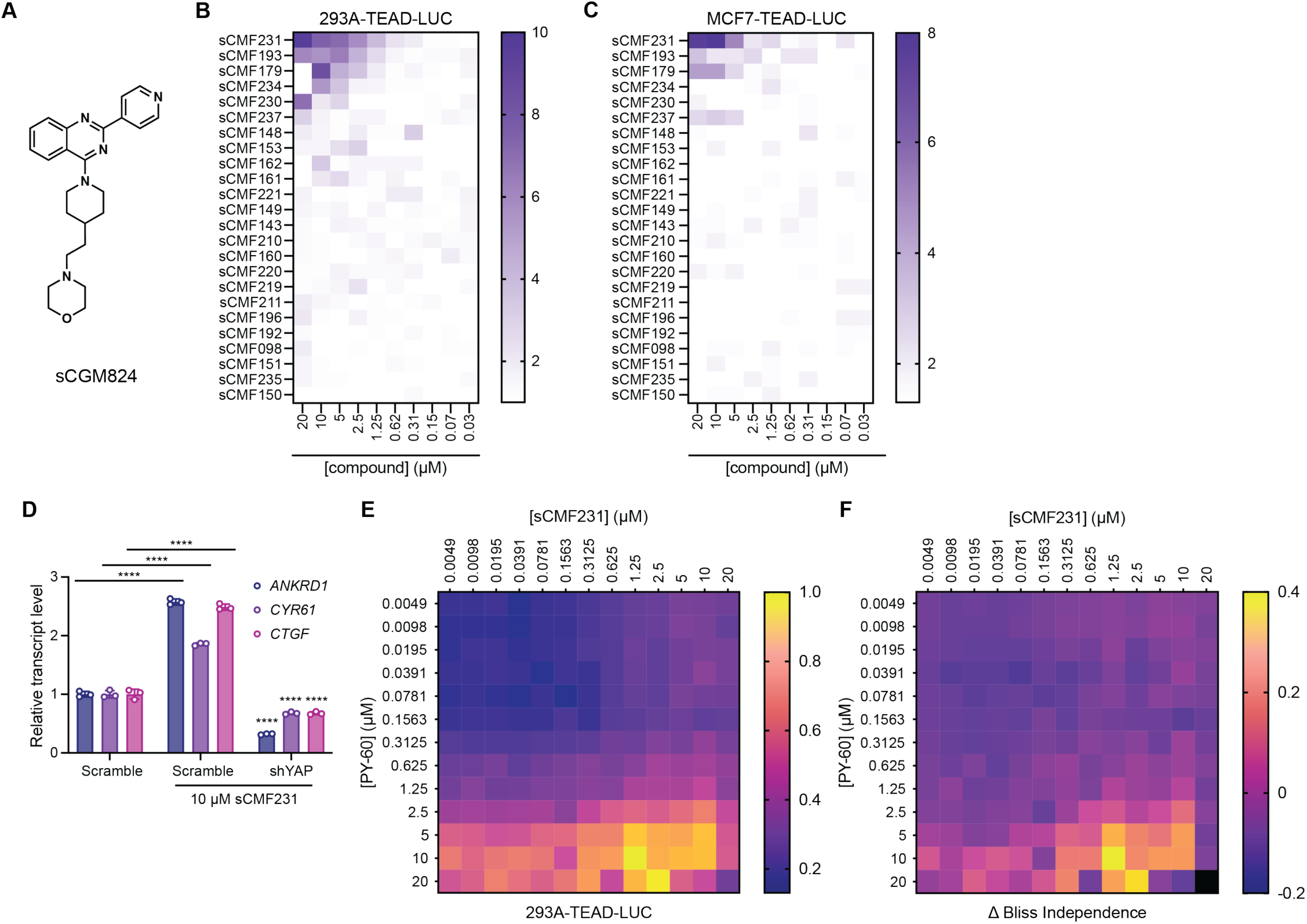
Discovery of a quinazoline-based YAP activating chemical scaffold. (A) Structure of screening hit sCGM824. (B, C) Heat maps of activity of compounds from the structure-activity relationship study in (B) 293A-TEAD-LUC and (C) MCF7-TEAD-LUC cells. Darker colored boxes indicate higher relative luminance values. (D) Relative transcript levels of *ANKRD1, CYR61,* and *CTGF* from HEK293A cells expressing the indicated YAP targeting shRNA and then treated with 10 µM sCMF231, measured by RT-qPCR (n=3, mean and s.d.). Statistical significance was determined using one-way ANOVA per transcript (****p<0.0001). Asterisks above the sample treated with shRNA to YAP and sCMF231 refer to statistical significance as compared to scramble treated with sCMF231. (E) Additive effects and (F) Bliss independence calculations for 293A-TEAD-LUC cells cotreated with sCMF231 and PY-60.

**Figure S2.**
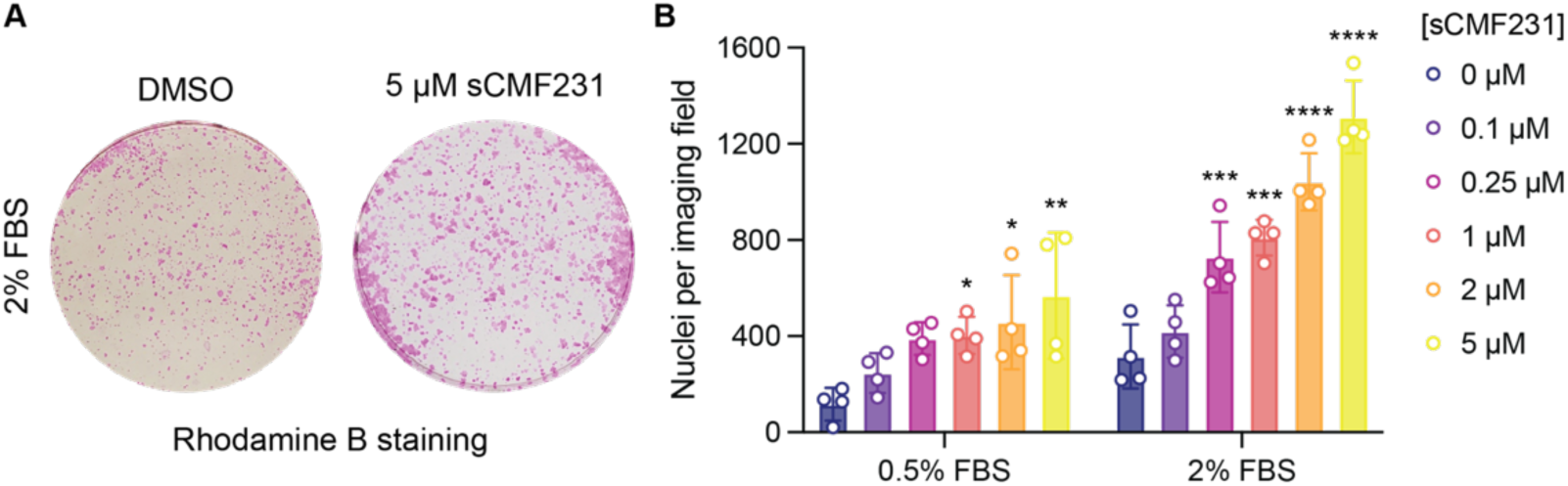
sCMF231 induces YAP-dependent proliferation. (A) Representative Rhodamine B staining, and (B) number of cells of human keratinocytes (HaCaT) in 0.5% or 2% FBS treated with sCMF231 for 7 days (n=4, mean and s.d.). Statistical analysis is a one-way ANOVA per serum condition (*p<0.05, **p<0.01, ***p<0.001, ****p<0.0001).

**Figure S3.**
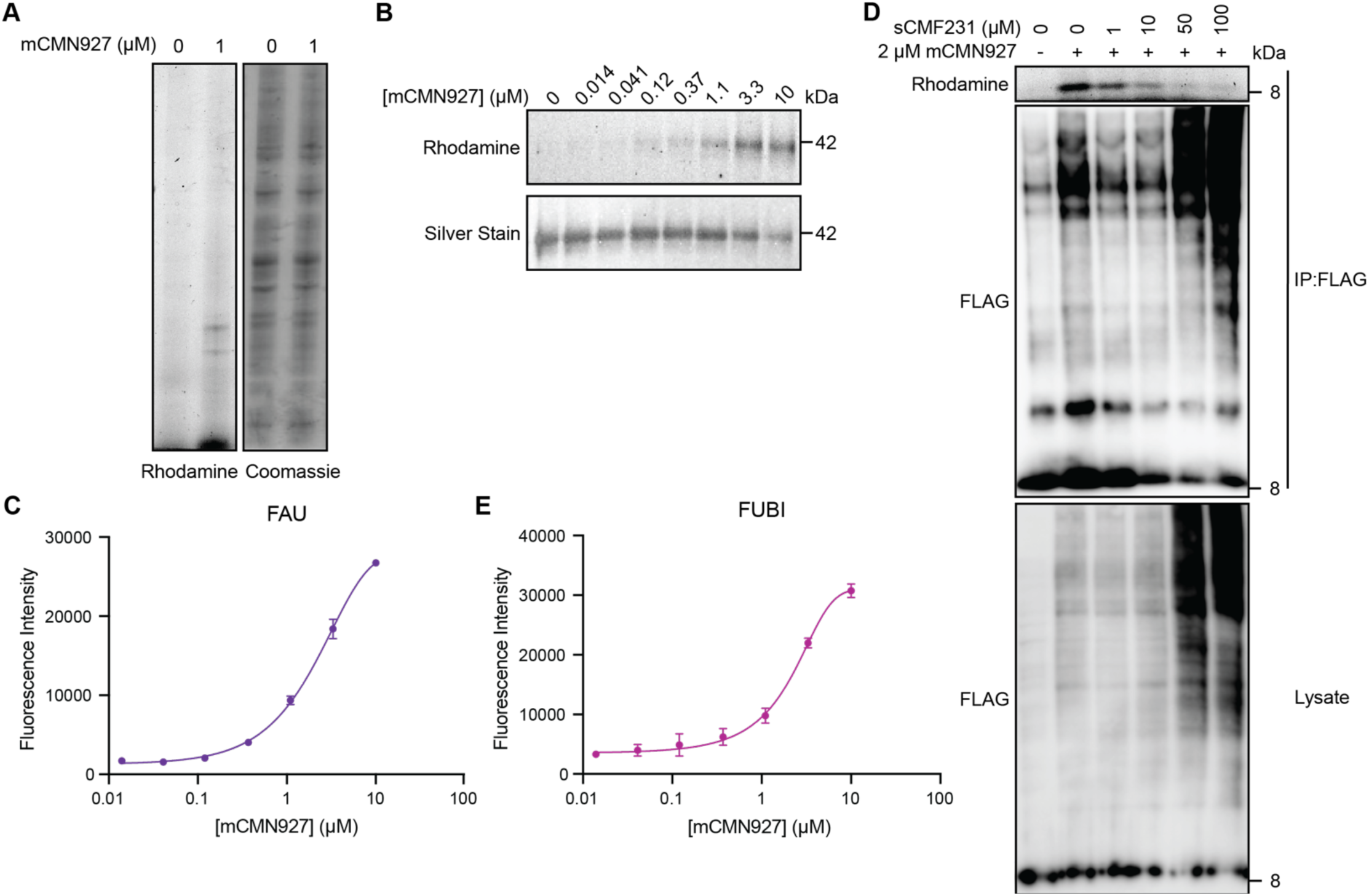
Chemical proteomics identifies FUBI as a druggable component of the Hippo pathway. (A) Representative fluorescent gel scans of rhodamine-azide labeled 293A cell lysates after *in situ* crosslinking with mCMN927. (B, C) Representative fluorescent gel scan and quantification of rhodamine azide-based labeling of recombinant GST-FAU exposed to mCMN927. (D) Representative fluorescent gel scan and anti-FLAG Western blot analysis of rhodamine-azide labeled material after FLAG-IP of FLAG-FUBI with treatment of 2 µM mCMN927 and competition with excess free sCMF231. (E) Quantification of rhodamine azide-based labeling of recombinant GST-FUBI exposed to mCMN927.

**Figure S4.**
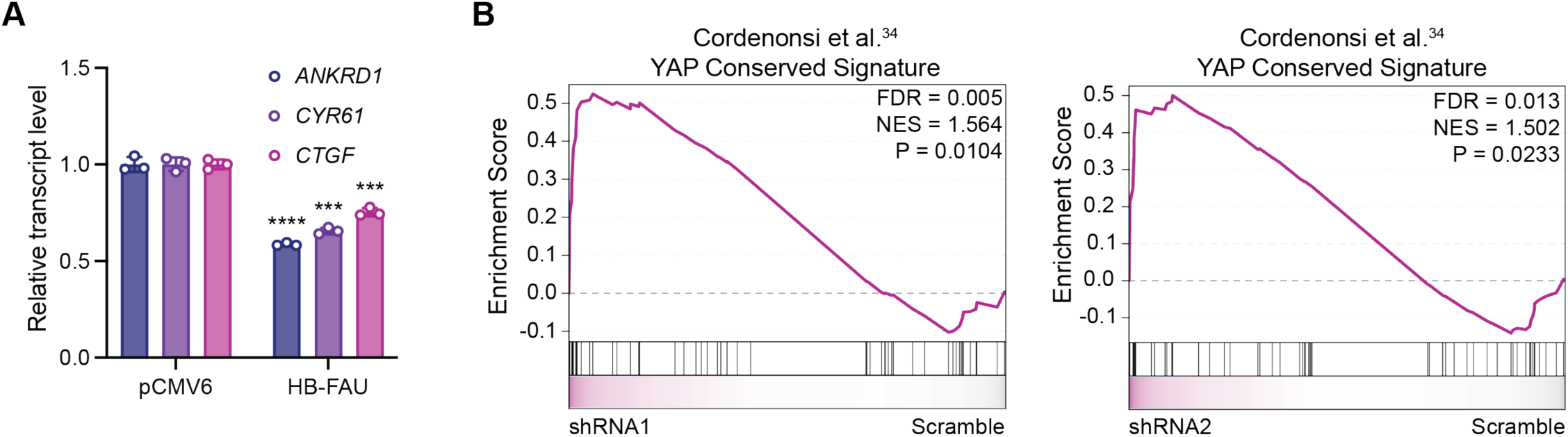
FAU regulates YAP activity. (A) Relative levels of YAP-dependent transcripts *ANKRD1, CYR61,* and *CTGF* from 293A cells transiently overexpressing FAU (n=3, mean and s.d.). Statistical significance was determined by univariate two-sided t-tests (***p<0.001, ****p<0.0001). (B) GSEA plots of a YAP-dependent curated gene sets (Cordenonsi et al.^34^ (MSigDB: M2871)) from 293A cells expressing FAU shRNA1 (left) or FAU shRNA2 (right).

**Figure S5.**
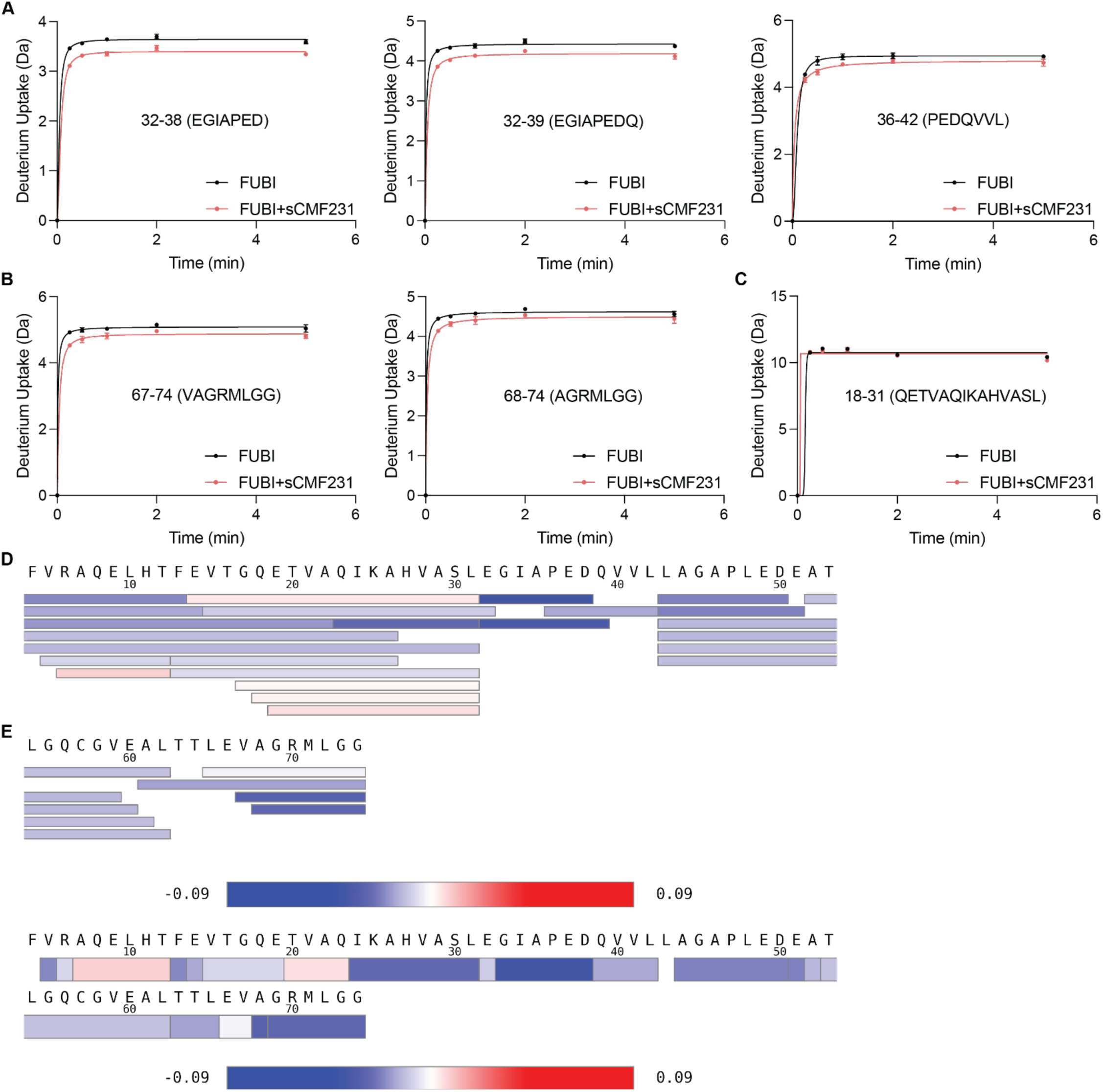
HDX-MS defines a potential binding site of sCMF231. (A, B) Traces of hydrogen/deuterium exchange as a function of deuteration time for peptides showing a reduction in solvent exchange occurring in FUBI upon incubation with 600 µM sCMF231. Peptides that displayed the largest decreases in solvent exchange upon sCMF231 binding were (A) residues 32-42 (EGIAPEDQVVL) and (B) residues 67-74 (VAGRMLGG). (C) Trace of hydrogen/deuterium exchange as a function of deuteration time for one example peptide showing no reduction in solvent exchange occurring in FUBI upon incubation with 600 µM sCMF231. (D) Coverage map and (E) heat map of all the peptides showing sequence coverage and redundancy of peptides that cover each amino acid. Blue on the color scale indicates decreased deuterium uptake and red on the color scale indicates increased deuterium uptake. (A-C) The averages are shown from three technical replicates, with error bars representing standard deviation.

**Figure S6.**
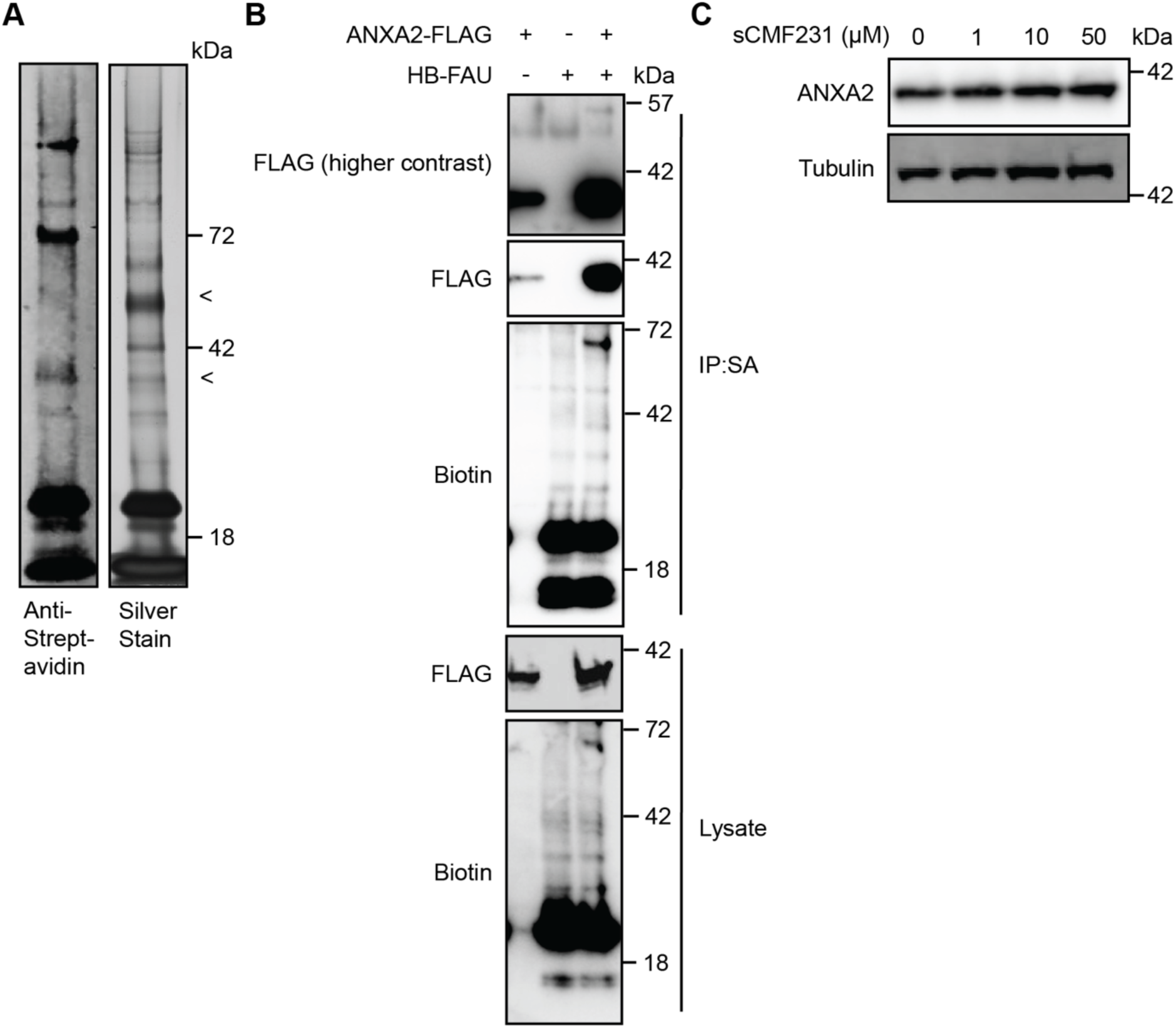
ANXA2 is a FUBI substrate. (A) Anti-Streptavidin Western blot and silver stained gel of streptavidin-enriched material after purification of HB-FUBI by Ni-NTA chromatography. Arrows indicate bands pooled and analyzed by LC-MS/MS containing ANXA2. (B) Immunoblotting analysis of ANXA2-FLAG from streptavidin-enriched content from HEK293T cells. (C) Anti-ANXA2 Western blot of HEK293A cells treated with the indicated concentrations of sCMF231 for 24h.

**Figure S7.**
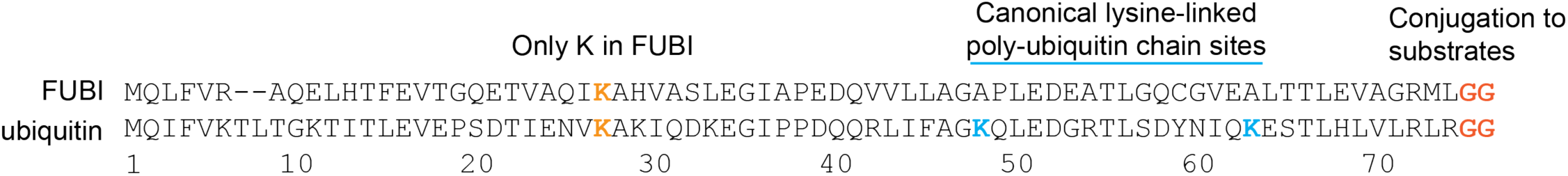
Amino acid sequences of FUBI and ubiquitin. The only lysine in FUBI is K25. FUBI contains the C-terminal -GG required for conjugation to target substrates.

**Figure S8.**
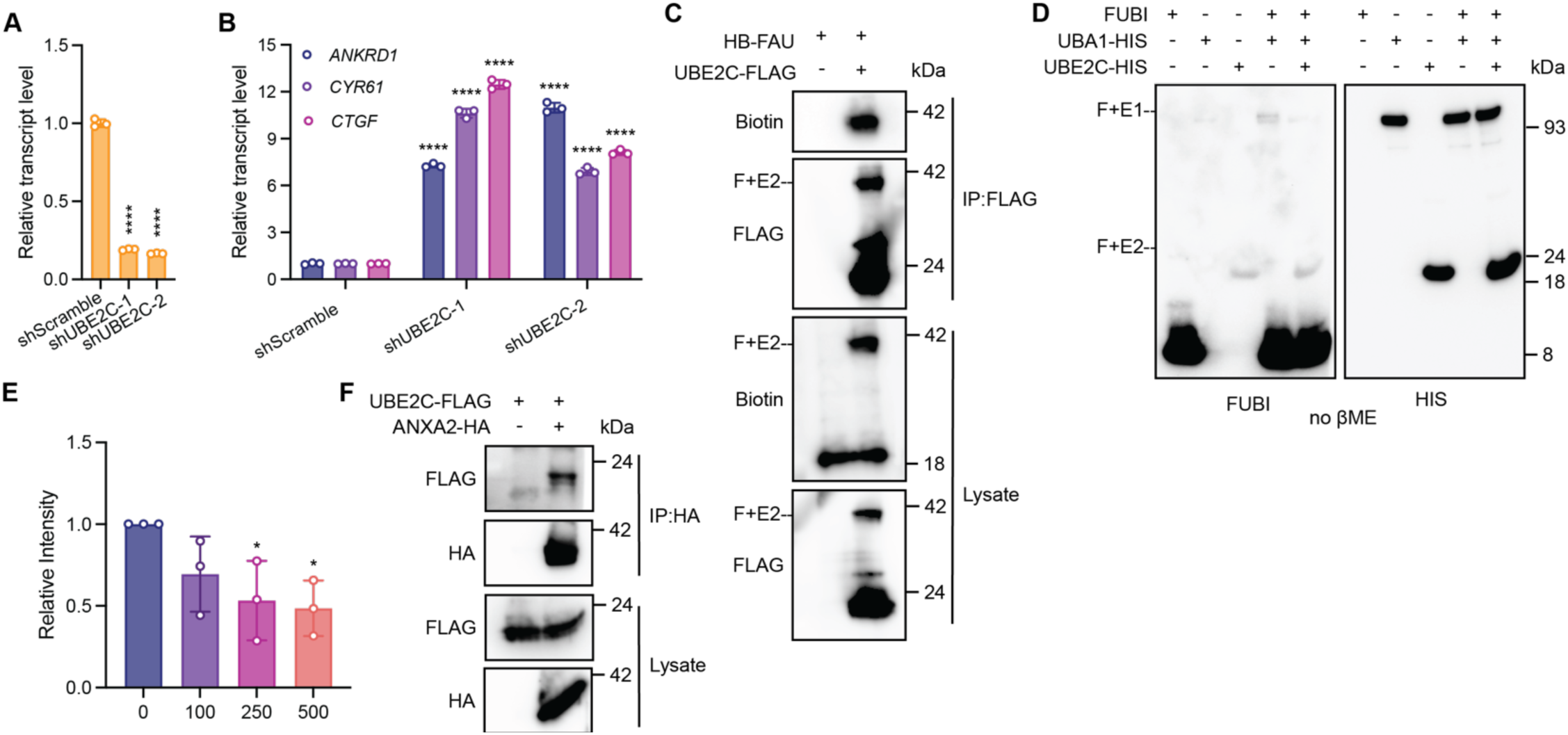
UBE2C plays an integral role in ANXA2 fubylation. (A) Relative transcript levels of UBE2C following knockdown by the indicated shRNA, measured by RT-qPCR (n=3, mean and s.d.). (B) Relative transcript levels of YAP-dependent genes (*ANKRD1, CTGF, CYR61*) expressing the indicated shRNA, measured by RT-qPCR (n=3, mean and s.d.). (C) Immunoblotting analysis of HB-FAU from anti-FLAG immunoprecipitated content from HEK293T cells. (D) Anti-FUBI Western blot showing the interaction of UBE2C and FUBI *in vitro* is competed by βME. (E) Quantification of Figure 5D indicating sCMF231 decreases the transfer of FUBI from UBA1 to UBE2C. (F) Immunoblotting analysis of UBE2C-FLAG from anti-HA immunoprecipitated content from HEK293T cells. Statistical analyses (A,B,E) are one-way ANOVA (*p<0.05, ****p<0.0001). (C,D) F+E1 indicates the covalent complex of FUBI with UBA1. F+E2 indicates the covalent complex of FUBI with UBE2C.

**Figure S9.**
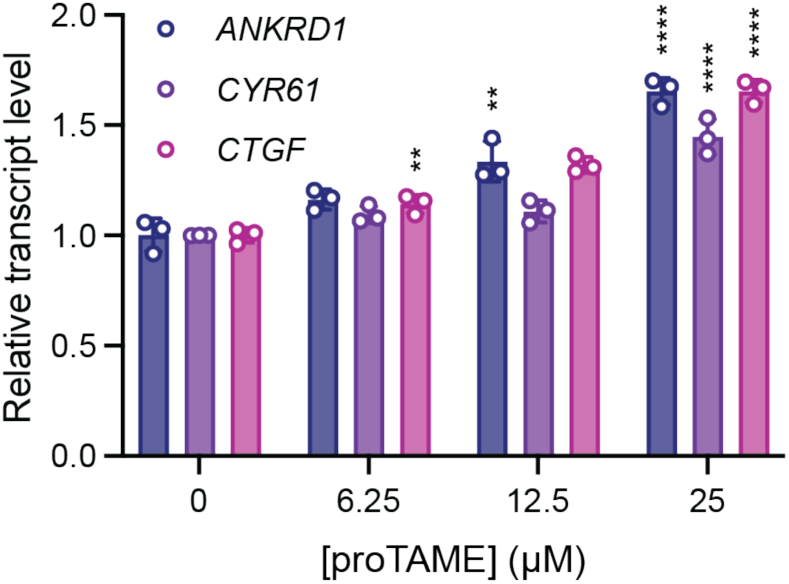
Inhibition of APC/C by proTAME moderately induces YAP-controlled transcripts. (A) Relative transcript levels of YAP-dependent genes (*ANKRD1, CYR61,* and *CTGF*) from HEK293A cells treated for 24h with proTAME, measured by RT-qPCR (n=3, mean and s.d.). Statistical analysis is one-way ANOVA (**p<0.01, ****p<0.0001).

## References

(1) Xu, T.; Wang, W.; Zhang, S.; Stewart, R. A.; Yu, W. Identifying Tumor Suppressors in Genetic Mosaics: The Drosophila Lats Gene Encodes a Putative Protein Kinase. Development 1995, 121 (4), 1053–1063. 10.1242/dev.121.4.1053.

(2) Wu, S.; Huang, J.; Dong, J.; Pan, D. Hippo Encodes a Ste-20 Family Protein Kinase That Restricts Cell Proliferation and Promotes Apoptosis in Conjunction with Salvador and Warts. Cell 2003, 114 (4), 445–456. 10.1016/S0092-8674(03)00549-X.

(3) Zhao, B.; Wei, X.; Li, W.; Udan, R. S.; Yang, Q.; Kim, J.; Xie, J.; Ikenoue, T.; Yu, J.; Li, L.; Zheng, P.; Ye, K.; Chinnaiyan, A.; Halder, G.; Lai, Z.-C.; Guan, K.-L. Inactivation of YAP Oncoprotein by the Hippo Pathway Is Involved in Cell Contact Inhibition and Tissue Growth Control. Genes Dev. 2007, 21 (21), 2747–2761. 10.1101/gad.1602907.

(4) Zhao, B.; Li, L.; Tumaneng, K.; Wang, C.-Y.; Guan, K.-L. A Coordinated Phosphorylation by Lats and CK1 Regulates YAP Stability through SCFβ-TRCP. Genes Dev. 2010, 24 (1), 72–85. 10.1101/gad.1843810.

(5) Huang, J.; Wu, S.; Barrera, J.; Matthews, K.; Pan, D. The Hippo Signaling Pathway Coordinately Regulates Cell Proliferation and Apoptosis by Inactivating Yorkie, the Drosophila Homolog of YAP. Cell 2005, 122 (3), 421–434. 10.1016/j.cell.2005.06.007.

(6) Chan, E. H. Y.; Nousiainen, M.; Chalamalasetty, R. B.; Schäfer, A.; Nigg, E. A.; Silljé, H. H. W. The Ste20-like Kinase Mst2 Activates the Human Large Tumor Suppressor Kinase Lats1. Oncogene 2005, 24 (12), 2076–2086. 10.1038/sj.onc.1208445.

(7) Zhao, B.; Ye, X.; Yu, J.; Li, L.; Li, W.; Li, S.; Yu, J.; Lin, J. D.; Wang, C.-Y.; Chinnaiyan, A. M.; Lai, Z.-C.; Guan, K.-L. TEAD Mediates YAP-Dependent Gene Induction and Growth Control. Genes Dev. 2008, 22 (14), 1962–1971. 10.1101/gad.1664408.

(8) Lai, Z.-C.; Wei, X.; Shimizu, T.; Ramos, E.; Rohrbaugh, M.; Nikolaidis, N.; Ho, L.-L.; Li, Y. Control of Cell Proliferation and Apoptosis by Mob as Tumor Suppressor, Mats. Cell 2005, 120 (5), 675–685. 10.1016/j.cell.2004.12.036.

(9) Su, T.; Ludwig, M. Z.; Xu, J.; Fehon, R. G. Kibra and Merlin Activate the Hippo Pathway Spatially Distinct from and Independent of Expanded. Dev. Cell 2017, 40 (5), 478–490.e3. 10.1016/j.devcel.2017.02.004.

(10) Harvey, K. F.; Pfleger, C. M.; Hariharan, I. K. The *Drosophila* Mst Ortholog, *Hippo*, Restricts Growth and Cell Proliferation and Promotes Apoptosis. Cell 2003, 114 (4), 457–467. 10.1016/S0092-8674(03)00557-9.

(11) Hergovich, A.; Schmitz, D.; Hemmings, B. A. The Human Tumour Suppressor LATS1 Is Activated by Human MOB1 at the Membrane. Biochem. Biophys. Res. Commun. 2006, 345 (1), 50–58. 10.1016/j.bbrc.2006.03.244.

(12) Yin, F.; Yu, J.; Zheng, Y.; Chen, Q.; Zhang, N.; Pan, D. Spatial Organization of Hippo Signaling at the Plasma Membrane Mediated by the Tumor Suppressor Merlin/NF2. Cell 2013, 154 (6), 1342–1355. 10.1016/j.cell.2013.08.025.

(13) Grieve, A. G.; Moss, S. E.; Hayes, M. J. Annexin A2 at the Interface of Actin and Membrane Dynamics: A Focus on Its Roles in Endocytosis and Cell Polarization. Int. J. Cell Biol. 2012, 2012, 852430. 10.1155/2012/852430.

(14) Lee, D. B. N.; Jamgotchian, N.; Allen, S. G.; Kan, F. W. K.; Hale, I. L. Annexin A2 Heterotetramer: Role in Tight Junction Assembly. Am. J. Physiol. Renal Physiol. 2004, 287 (3), F481–491. 10.1152/ajprenal.00175.2003.

(15) Shalhout, S. Z.; Yang, P.-Y.; Grzelak, E. M.; Nutsch, K.; Shao, S.; Zambaldo, C.; Iaconelli, J.; Ibrahim, L.; Stanton, C.; Chadwick, S. R.; Chen, E.; DeRan, M.; Li, S.; Hull, M.; Wu, X.; Chatterjee, A. K.; Shen, W.; Camargo, F. D.; Schultz, P. G.; Bollong, M. J. YAP-Dependent Proliferation by a Small Molecule Targeting Annexin A2. Nat. Chem. Biol. 2021, 17 (7), 767–775. 10.1038/s41589-021-00755-0.

(16) Moya, I. M.; Halder, G. Hippo–YAP/TAZ Signalling in Organ Regeneration and Regenerative Medicine. Nat. Rev. Mol. Cell Biol. 2019, 20 (4), 211–226. 10.1038/s41580-018-0086-y.

(17) Cai, J.; Zhang, N.; Zheng, Y.; de Wilde, R. F.; Maitra, A.; Pan, D. The Hippo Signaling Pathway Restricts the Oncogenic Potential of an Intestinal Regeneration Program. Genes Dev. 2010, 24 (21), 2383–2388. 10.1101/gad.1978810.

(18) Elbediwy, A.; Vincent-Mistiaen, Z. I.; Spencer-Dene, B.; Stone, R. K.; Boeing, S.; Wculek, S. K.; Cordero, J.; Tan, E. H.; Ridgway, R.; Brunton, V. G.; Sahai, E.; Gerhardt, H.; Behrens, A.; Malanchi, I.; Sansom, O. J.; Thompson, B. J. Integrin Signalling Regulates YAP and TAZ to Control Skin Homeostasis. Dev. Camb. Engl. 2016, 143 (10), 1674–1687. 10.1242/dev.133728.

(19) Grijalva, J. L.; Huizenga, M.; Mueller, K.; Rodriguez, S.; Brazzo, J.; Camargo, F.; Sadri-Vakili, G.; Vakili, K. Dynamic Alterations in Hippo Signaling Pathway and YAP Activation during Liver Regeneration. Am. J. Physiol.-Gastrointest. Liver Physiol. 2014, 307 (2), G196–G204. 10.1152/ajpgi.00077.2014.

(20) Nowell, C. S.; Odermatt, P. D.; Azzolin, L.; Hohnel, S.; Wagner, E. F.; Fantner, G. E.; Lutolf, M. P.; Barrandon, Y.; Piccolo, S.; Radtke, F. Chronic Inflammation Imposes Aberrant Cell Fate in Regenerating Epithelia through Mechanotransduction. Nat. Cell Biol. 2016, 18 (2), 168–180. 10.1038/ncb3290.

(21) Deng, Y.; Wu, A.; Li, P.; Li, G.; Qin, L.; Song, H.; Mak, K. K. Yap1 Regulates Multiple Steps of Chondrocyte Differentiation during Skeletal Development and Bone Repair. Cell Rep. 2016, 14 (9), 2224– 2237. 10.1016/j.celrep.2016.02.021.

(22) Loforese, G.; Malinka, T.; Keogh, A.; Baier, F.; Simillion, C.; Montani, M.; Halazonetis, T. D.; Candinas, D.; Stroka, D. Impaired Liver Regeneration in Aged Mice Can Be Rescued by Silencing Hippo Core Kinases MST1 and MST2. EMBO Mol. Med. 2017, 9 (1), 46–60. 10.15252/emmm.201506089.

(23) Heallen, T.; Morikawa, Y.; Leach, J.; Tao, G.; Willerson, J. T.; Johnson, R. L.; Martin, J. F. Hippo Signaling Impedes Adult Heart Regeneration. Development 2013, 140 (23), 4683–4690. 10.1242/dev.102798.

(24) Leach, J. P.; Heallen, T.; Zhang, M.; Rahmani, M.; Morikawa, Y.; Hill, M. C.; Segura, A.; Willerson, J. T.; Martin, J. F. Hippo Pathway Deficiency Reverses Systolic Heart Failure after Infarction. Nature 2017, 550 (7675), 260–264. 10.1038/nature24045.

(25) Liu, S.; Li, K.; Wagner Florencio, L.; Tang, L.; Heallen, T. R.; Leach, J. P.; Wang, Y.; Grisanti, F.; Willerson, J. T.; Perin, E. C.; Zhang, S.; Martin, J. F. Gene Therapy Knockdown of Hippo Signaling Induces Cardiomyocyte Renewal in Pigs after Myocardial Infarction. Sci. Transl. Med. 2021, 13 (600), eabd6892. 10.1126/scitranslmed.abd6892.

(26) Grzelak, E. M.; Elshan, N. G. R. D.; Shao, S.; Bulos, M. L.; Joseph, S. B.; Chatterjee, A. K.; Chen, J. J.; Nguyên-Trân, V.; Schultz, P. G.; Bollong, M. J. Pharmacological YAP Activation Promotes Regenerative Repair of Cutaneous Wounds. Proc. Natl. Acad. Sci. 2023, 120 (28), e2305085120. 10.1073/pnas.2305085120.

(27) Fan, F.; He, Z.; Kong, L.-L.; Chen, Q.; Yuan, Q.; Zhang, S.; Ye, J.; Liu, H.; Sun, X.; Geng, J.; Yuan, L.; Hong, L.; Xiao, C.; Zhang, W.; Sun, X.; Li, Y.; Wang, P.; Huang, L.; Wu, X.; Ji, Z.; Wu, Q.; Xia, N.-S.; Gray, N. S.; Chen, L.; Yun, C.-H.; Deng, X.; Zhou, D. Pharmacological Targeting of Kinases MST1 and MST2 Augments Tissue Repair and Regeneration. Sci. Transl. Med. 2016, 8 (352). 10.1126/scitranslmed.aaf2304.

(28) Namoto, K.; Baader, C.; Orsini, V.; Landshammer, A.; Breuer, E.; Dinh, K. T.; Ungricht, R.; Pikiolek, M.; Laurent, S.; Lu, B.; Aebi, A.; Schönberger, K.; Vangrevelinghe, E.; Evrova, O.; Sun, T.; Annunziato, S.; Lachal, J.; Redmond, E.; Wang, L.; Wetzel, K.; Capodieci, P.; Turner, J.; Schutzius, G.; Unterreiner, V.; Trunzer, M.; Buschmann, N.; Behnke, D.; Machauer, R.; Scheufler, C.; Parker, C. N.; Ferro, M.; Grevot, A.; Beyerbach, A.; Lu, W.-Y.; Forbes, S. J.; Wagner, J.; Bouwmeester, T.; Liu, J.; Sohal, B.; Sahambi, S.; Greenbaum, L. E.; Lohmann, F.; Hoppe, P.; Cong, F.; Sailer, A. W.; Ruffner, H.; Glatthar, R.; Humar, B.; Clavien, P.-A.; Dill, M. T.; George, E.; Maibaum, J.; Liberali, P.; Tchorz, J. S. NIBR-LTSi Is a Selective LATS Kinase Inhibitor Activating YAP Signaling and Expanding Tissue Stem Cells *in Vitro* and *in Vivo*. Cell Stem Cell 2024, 31 (4), 554–569.e17. 10.1016/j.stem.2024.03.003.

(29) Li, T.; Wen, Yiqiong; Lu, Qiongfen; Hua, Shu; Hou, Yunjiao; Du, Xiaohua; Zheng, Yuanyuan; and Sun, S. MST1/2 in Inflammation and Immunity. Cell Adhes. Migr. 2023, 17 (1), 1–15. 10.1080/19336918.2023.2276616.

(30) Zhu, Y.; Abedini, A.; Rodriguez, G. M.; McCloskey, C. W.; Abou-Hamad, J.; Salah, O. S.; Larocque, J.; Tsoi, M. F.; Boerboom, D.; Cook, D.; Vanderhyden, B. Loss of LATS1 and LATS2 Promotes Ovarian Tumor Formation by Enhancing AKT Activity and PD-L1 Expression. Oncogene 2025, 1–13. 10.1038/s41388-025-03387-z.

(31) Galan, J. A.; Avruch, J. MST1/MST2 Protein Kinases: Regulation and Physiologic Roles. Biochemistry 2016, 55 (39), 5507–5519. 10.1021/acs.biochem.6b00763.

(32) Hergovich, A.; Hemmings, B. A. Hippo Signalling in the G2/M Cell Cycle Phase: Lessons Learned from the Yeast MEN and SIN Pathways. Semin. Cell Dev. Biol. 2012, 23 (7), 794–802. 10.1016/j.semcdb.2012.04.001.

(33) Praskova, M.; Xia, F.; Avruch, J. MOBKL1A/MOBKL1B Phosphorylation by MST1 and MST2 Inhibits Cell Proliferation. Curr. Biol. 2008, 18 (5), 311–321. 10.1016/j.cub.2008.02.006.

(34) Cordenonsi, M.; Zanconato, F.; Azzolin, L.; Forcato, M.; Rosato, A.; Frasson, C.; Inui, M.; Montagner, M.; Parenti, A. R.; Poletti, A.; Daidone, M. G.; Dupont, S.; Basso, G.; Bicciato, S.; Piccolo, S. The Hippo Transducer TAZ Confers Cancer Stem Cell-Related Traits on Breast Cancer Cells. Cell 2011, 147 (4), 759–772. 10.1016/j.cell.2011.09.048.

(35) Bulos, M. L.; Grzelak, E. M.; Li-Ma, C.; Chen, E.; Hull, M.; Johnson, K. A.; Bollong, M. J. Pharmacological Inhibition of CLK2 Activates YAP by Promoting Alternative Splicing of AMOTL2. eLife 2023, 12, RP88508. 10.7554/eLife.88508.

(36) Zhao, W.; Sachsenmeier, K.; Zhang, L.; Sult, E.; Hollingsworth, R. E.; Yang, H. A New Bliss Independence Model to Analyze Drug Combination Data. SLAS Discov. 2014, 19 (5), 817–821. 10.1177/1087057114521867.

(37) van den Heuvel, J.; Ashiono, C.; Gillet, L. C.; Dörner, K.; Wyler, E.; Zemp, I.; Kutay, U. Processing of the Ribosomal Ubiquitin-like Fusion Protein FUBI-eS30/FAU Is Required for 40S Maturation and Depends on USP36. eLife 2021, 10, e70560. 10.7554/eLife.70560.

(38) O’Dea, R.; Kazi, N.; Hoffmann-Benito, A.; Zhao, Z.; Recknagel, S.; Wendrich, K.; Janning, P.; Gersch, M. Molecular Basis for Ubiquitin/Fubi Cross-Reactivity in USP16 and USP36. Nat. Chem. Biol. 2023, 19 (11), 1394–1405. 10.1038/s41589-023-01388-1.

(39) Kas, K.; Michiels, L.; Merregaert, J. Genomic Structure and Expression of the Human Fau Gene: Encoding the Ribosomal Protein S30 Fused to a Ubiquitin-like Protein. Biochem. Biophys. Res. Commun. 1992, 187 (2), 927–933. 10.1016/0006-291X(92)91286-Y.

(40) Ozohanics, O.; Ambrus, A. Hydrogen-Deuterium Exchange Mass Spectrometry: A Novel Structural Biology Approach to Structure, Dynamics and Interactions of Proteins and Their Complexes. Life 2020, 10 (11), 286. 10.3390/life10110286.

(41) Nakamura, M.; Xavier, R. M.; Tsunematsu, T.; Tanigawa, Y. Molecular Cloning and Characterization of a cDNA Encoding Monoclonal Nonspecific Suppressor Factor. Proc. Natl. Acad. Sci. 1995, 92 (8), 3463–3467. 10.1073/pnas.92.8.3463.

(42) Nakamura, M.; Watanabe, N.; Notsu, K. Ubiquitin-like Protein MNSFβ Covalently Binds to Cytosolic 10-Formyltetrahydrofolate Dehydrogenase and Regulates Thymocyte Function. Biochem. Biophys. Res. Commun. 2015, 464 (4), 1096–1100. 10.1016/j.bbrc.2015.07.083.

(43) Nakamura, M.; Tanigawa, Y. Characterization of Ubiquitin-like Polypeptide Acceptor Protein, a Novel pro-Apoptotic Member of the Bcl2 Family: Ubiquitin-like Polypeptide Acceptor Protein. Eur. J. Biochem. 2003, 270 (20), 4052–4058. 10.1046/j.1432-1033.2003.03790.x.

(44) Nakamura, M.; Shimosaki, S. The Ubiquitin-like Protein Monoclonal Nonspecific Suppressor Factor β Conjugates to Endophilin II and Regulates Phagocytosis: MNSFβ Conjugates to Endophilin II. FEBS J. 2009, 276 (21), 6355–6363. 10.1111/j.1742-4658.2009.07348.x.

(45) Tagwerker, C.; Flick, K.; Cui, M.; Guerrero, C.; Dou, Y.; Auer, B.; Baldi, P.; Huang, L.; Kaiser, P. A Tandem Affinity Tag for Two-Step Purification under Fully Denaturing Conditions: Application in Ubiquitin Profiling and Protein Complex Identification Combined with in vivoCross-Linking*. Mol. Cell. Proteomics 2006, 5 (4), 737–748. 10.1074/mcp.M500368-MCP200.

(46) Castañeda, C. A.; Dixon, E. K.; Walker, O.; Chaturvedi, A.; Nakasone, M. A.; Curtis, J. E.; Reed, M. R.; Krueger, S.; Cropp, T. A.; Fushman, D. Linkage via K27 Bestows Ubiquitin Chains with Unique Properties among Polyubiquitins. Structure 2016, 24 (3), 423–436. 10.1016/j.str.2016.01.007.

(47) Nakagawa, T.; Nakayama, K. Protein Monoubiquitylation: Targets and Diverse Functions. Genes Cells 2015, 20 (7), 543–562. 10.1111/gtc.12250.

(48) Komander, D.; Rape, M. The Ubiquitin Code. Annu. Rev. Biochem. 2012, 81 (1), 203–229. 10.1146/annurev-biochem-060310-170328.

(49) Cappadocia, L.; Lima, C. D. Ubiquitin-like Protein Conjugation: Structures, Chemistry, and Mechanism. Chem. Rev. 2018, 118 (3), 889–918. 10.1021/acs.chemrev.6b00737.

(50) Schulman, B. A.; Wade Harper, J. Ubiquitin-like Protein Activation by E1 Enzymes: The Apex for Downstream Signalling Pathways. Nat. Rev. Mol. Cell Biol. 2009, 10 (5), 319–331. 10.1038/nrm2673.

(51) Rotin, D.; Kumar, S. Physiological Functions of the HECT Family of Ubiquitin Ligases. Nat. Rev. Mol. Cell Biol. 2009, 10 (6), 398–409. 10.1038/nrm2690.

(52) Deshaies, R. J.; Joazeiro, C. A. P. RING Domain E3 Ubiquitin Ligases. Annu. Rev. Biochem. 2009, 78 (Volume 78, 2009), 399–434. 10.1146/annurev.biochem.78.101807.093809.

(53) Yamano, H. APC/C: Current Understanding and Future Perspectives. F1000Research 2019, 8, F1000 Faculty Rev-725. 10.12688/f1000research.18582.1.

(54) Tang, Z.; Li, B.; Bharadwaj, R.; Zhu, H.; Özkan, E.; Hakala, K.; Deisenhofer, J.; Yu, H. APC2 Cullin Protein and APC11 RING Protein Comprise the Minimal Ubiquitin Ligase Module of the Anaphase-Promoting Complex. Mol. Biol. Cell 2001, 12 (12), 3839–3851. 10.1091/mbc.12.12.3839.

(55) Zeng, X.; Sigoillot, F.; Gaur, S.; Choi, S.; Pfaff, K. L.; Oh, D.-C.; Hathaway, N.; Dimova, N.; Cuny, G. D.; King, R. W. Pharmacologic Inhibition of the Anaphase-Promoting Complex Induces A Spindle Checkpoint-Dependent Mitotic Arrest in the Absence of Spindle Damage. Cancer Cell 2010, 18 (4), 382–395. 10.1016/j.ccr.2010.08.010.

(56) Hyer, M. L.; Milhollen, M. A.; Ciavarri, J.; Fleming, P.; Traore, T.; Sappal, D.; Huck, J.; Shi, J.; Gavin, J.; Brownell, J.; Yang, Y.; Stringer, B.; Griffin, R.; Bruzzese, F.; Soucy, T.; Duffy, J.; Rabino, C.; Riceberg, J.; Hoar, K.; Lublinsky, A.; Menon, S.; Sintchak, M.; Bump, N.; Pulukuri, S. M.; Langston, S.; Tirrell, S.; Kuranda, M.; Veiby, P.; Newcomb, J.; Li, P.; Wu, J. T.; Powe, J.; Dick, L. R.; Greenspan, P.; Galvin, K.; Manfredi, M.; Claiborne, C.; Amidon, B. S.; Bence, N. F. A Small-Molecule Inhibitor of the Ubiquitin Activating Enzyme for Cancer Treatment. Nat. Med. 2018, 24 (2), 186–193. 10.1038/nm.4474.

(57) Langston, S. P.; Grossman, S.; England, D.; Afroze, R.; Bence, N.; Bowman, D.; Bump, N.; Chau, R.; Chuang, B.-C.; Claiborne, C.; Cohen, L.; Connolly, K.; Duffey, M.; Durvasula, N.; Freeze, S.; Gallery, M.; Galvin, K.; Gaulin, J.; Gershman, R.; Greenspan, P.; Grieves, J.; Guo, J.; Gulavita, N.; Hailu, S.; He, X.; Hoar, K.; Hu, Y.; Hu, Z.; Ito, M.; Kim, M.-S.; Lane, S. W.; Lok, D.; Lublinsky, A.; Mallender, W.; McIntyre, C.; Minissale, J.; Mizutani, H.; Mizutani, M.; Molchinova, N.; Ono, K.; Patil, A.; Qian, M.; Riceberg, J.; Shindi, V.; Sintchak, M. D.; Song, K.; Soucy, T.; Wang, Y.; Xu, H.; Yang, X.; Zawadzka, A.; Zhang, J.; Pulukuri, S. M. Discovery of TAK-981, a First-in-Class Inhibitor of SUMO-Activating Enzyme for the Treatment of Cancer. J. Med. Chem. 2021, 64 (5), 2501–2520. 10.1021/acs.jmedchem.0c01491.

(58) Ceccarelli, D. F.; Tang, X.; Pelletier, B.; Orlicky, S.; Xie, W.; Plantevin, V.; Neculai, D.; Chou, Y.-C.; Ogunjimi, A.; Al-Hakim, A.; Varelas, X.; Koszela, J.; Wasney, G. A.; Vedadi, M.; Dhe-Paganon, S.; Cox, S.; Xu, S.; Lopez-Girona, A.; Mercurio, F.; Wrana, J.; Durocher, D.; Meloche, S.; Webb, D. R.; Tyers, M.; Sicheri, F. An Allosteric Inhibitor of the Human Cdc34 Ubiquitin-Conjugating Enzyme. Cell 2011, 145 (7), 1075–1087. 10.1016/j.cell.2011.05.039.

(59) LaPlante, G.; Zhang, W. Targeting the Ubiquitin-Proteasome System for Cancer Therapeutics by Small-Molecule Inhibitors. Cancers 2021, 13 (12), 3079. 10.3390/cancers13123079.

(60) Li, W.; Garcia-Rivera, E. M.; Mitchell, D. C.; Chick, J. M.; Maetani, M.; Knapp, J. M.; Matthews, G. M.; Shirasaki, R.; de Matos Simoes, R.; Viswanathan, V.; Pulice, J. L.; Rees, M. G.; Roth, J. A.; Gygi, S. P.; Mitsiades, C. S.; Kadoch, C.; Schreiber, S. L.; Ostrem, J. M. L. Highly Specific Intracellular Ubiquitination of a Small Molecule. BioRxiv Prepr. Serv. Biol. 2024, 2024.01.26.577493. 10.1101/2024.01.26.577493.

(61) Kochańczyk, T.; Hann, Z. S.; Lux, M. C.; Delos Reyes, A. M. V.; Ji, C.; Tan, D. S.; Lima, C. D. Structural Basis for Transthiolation Intermediates in the Ubiquitin Pathway. Nature 2024, 633 (8028), 216–223. 10.1038/s41586-024-07828-9.

(62) Lv, Z.; Williams, K. M.; Yuan, L.; Atkison, J. H.; Olsen, S. K. Crystal Structure of a Human Ubiquitin E1– Ubiquitin Complex Reveals Conserved Functional Elements Essential for Activity. J. Biol. Chem. 2018, 293 (47), 18337–18352. 10.1074/jbc.RA118.003975.

(63) Olsen, S. K.; Lima, C. D. Structure of a Ubiquitin E1-E2 Complex: Insights to E1-E2 Thioester Transfer. Mol. Cell 2013, 49 (5), 884–896. 10.1016/j.molcel.2013.01.013.

(64) Zhan, P.; Lu, Y.; Lu, J.; Cheng, Y.; Luo, C.; Yang, F.; Xi, W.; Wang, J.; Cen, X.; Wang, F.; Xie, C.; Yin, Z. The Activation of the Notch Signaling Pathway by UBE2C Promotes the Proliferation and Metastasis of Hepatocellular Carcinoma. Sci. Rep. 2024, 14 (1), 22859. 10.1038/s41598-024-72714-3.

(65) Zhang, S.; Shen, Y.; Li, H.; Bi, C.; Sun, Y.; Xiong, X.; Wei, W.; Sun, Y. The Negative Cross-Talk between SAG/RBX2/ROC2 and APC/C E3 Ligases in Regulation of Cell Cycle Progression and Drug Resistance. Cell Rep. 2020, 32 (10), 108102. 10.1016/j.celrep.2020.108102.

(66) Zhou, Z.; He, M.; Shah, A. A.; Wan, Y. Insights into APC/C: From Cellular Function to Diseases and Therapeutics. Cell Div. 2016, 11 (1), 9. 10.1186/s13008-016-0021-6.

(67) Kim, W.; Cho, Y. S.; Wang, X.; Park, O.; Ma, X.; Kim, H.; Gan, W.; Jho, E.; Cha, B.; Jeung, Y.; Zhang, L.; Gao, B.; Wei, W.; Jiang, J.; Chung, K.-S.; Yang, Y. Hippo Signaling Is Intrinsically Regulated during Cell Cycle Progression by APC/CCdh1. Proc. Natl. Acad. Sci. 2019, 116 (19), 9423–9432. 10.1073/pnas.1821370116.

(68) Gmachl, M.; Gieffers, C.; Podtelejnikov, A. V.; Mann, M.; Peters, J.-M. The RING-H2 Finger Protein APC11 and the E2 Enzyme UBC4 Are Sufficient to Ubiquitinate Substrates of the Anaphase-Promoting Complex. Proc. Natl. Acad. Sci. 2000, 97 (16), 8973–8978. 10.1073/pnas.97.16.8973.

(69) da Fonseca, P. C. A.; Kong, E. H.; Zhang, Z.; Schreiber, A.; Williams, M. A.; Morris, E. P.; Barford, D. Structures of APC/CCdh1 with Substrates Identify Cdh1 and Apc10 as the D-Box Co-Receptor. Nature 2011, 470 (7333), 274–278. 10.1038/nature09625.

(70) Liu, J.; Rothermund, C. A.; Ayala-Sanmartin, J.; Vishwanatha, J. K. Nuclear Annexin II Negatively Regulates Growth of LNCaP Cells and Substitution of Ser 11 and 25 to Glu Prevents Nucleo-Cytoplasmic Shuttling of Annexin II. BMC Biochem. 2003, 4 (1), 10. 10.1186/1471-2091-4-10.

(71) Jaquenoud, M.; van Drogen, F.; Peter, M. Cell Cycle-dependent Nuclear Export of Cdh1p May Contribute to the Inactivation of APC/CCdh1. EMBO J. 2002, 21 (23), 6515–6526. 10.1093/emboj/cdf634.

(72) Chan, A. H.; Lee, S. M. Y.; Chim, S. S.; Kok, L. D. S.; Waye, M. M. Y.; Lee, C.-Y.; Fung, K.-P.; Tsui, S. K. W. Molecular Cloning and Characterization of a RING-H2 Finger Protein, ANAPC11, the Human Homolog of Yeast Apc11p. J. Cell. Biochem. 2001, 83 (2), 249–258. 10.1002/jcb.1217.

(73) Garrod, D.; Chidgey, M. Desmosome Structure, Composition and Function. Biochim. Biophys. Acta BBA - Biomembr. 2008, 1778 (3), 572–587. 10.1016/j.bbamem.2007.07.014.

(74) Chen, S. N.; Gurha, P.; Lombardi, R.; Ruggiero, A.; Willerson, J. T.; Marian, A. The Hippo Pathway Is Activated and Is a Causal Mechanism for Adipogenesis in Arrhythmogenic Cardiomyopathy. Circ. Res. 2014, 114 (3), 454–468. 10.1161/CIRCRESAHA.114.302810.

(75) Giuliodori, A.; Beffagna, G.; Marchetto, G.; Fornetto, C.; Vanzi, F.; Toppo, S.; Facchinello, N.; Santimaria, M.; Vettori, A.; Rizzo, S.; Della Barbera, M.; Pilichou, K.; Argenton, F.; Thiene, G.; Tiso, N.; Basso, C. Loss of Cardiac Wnt/β-Catenin Signalling in Desmoplakin-Deficient AC8 Zebrafish Models Is Rescuable by Genetic and Pharmacological Intervention. Cardiovasc. Res. 2018, 114 (8), 1082–1097. 10.1093/cvr/cvy057.

(76) Dobin, A.; Davis, C. A.; Schlesinger, F.; Drenkow, J.; Zaleski, C.; Jha, S.; Batut, P.; Chaisson, M.; Gingeras, T. R. STAR: Ultrafast Universal RNA-Seq Aligner. Bioinformatics 2013, 29 (1), 15–21. 10.1093/bioinformatics/bts635.

(77) Liao, Y.; Smyth, G. K.; Shi, W. featureCounts: An Efficient General Purpose Program for Assigning Sequence Reads to Genomic Features. Bioinformatics 2014, 30 (7), 923–930. 10.1093/bioinformatics/btt656.

(78) Love, M. I.; Huber, W.; Anders, S. Moderated Estimation of Fold Change and Dispersion for RNA-Seq Data with DESeq2. Genome Biol. 2014, 15 (12), 550. 10.1186/s13059-014-0550-8.

(79) Subramanian, A.; Tamayo, P.; Mootha, V. K.; Mukherjee, S.; Ebert, B. L.; Gillette, M. A.; Paulovich, A.; Pomeroy, S. L.; Golub, T. R.; Lander, E. S.; Mesirov, J. P. Gene Set Enrichment Analysis: A Knowledge-Based Approach for Interpreting Genome-Wide Expression Profiles. Proc. Natl. Acad. Sci. 2005, 102 (43), 15545–15550. 10.1073/pnas.0506580102.

(80) Peacock, R. B.; Davis, J. R.; Markwick, P. R. L.; Komives, E. A. Dynamic Consequences of Mutation of Tryptophan 215 in Thrombin. Biochemistry 2018, 57 (18), 2694–2703. 10.1021/acs.biochem.8b00262.

(81) Wales, T. E.; Fadgen, K. E.; Gerhardt, G. C.; Engen, J. R. High-Speed and High-Resolution UPLC Separation at Zero Degrees Celsius. Anal. Chem. 2008, 80 (17), 6815–6820. 10.1021/ac8008862.

(82) Ramsey, K. M.; Dembinski, H. E.; Chen, W.; Ricci, C. G.; Komives, E. A. DNA and IκBα Both Induce Long-Range Conformational Changes in NFκB. J. Mol. Biol. 2017, 429 (7), 999–1008. 10.1016/j.jmb.2017.02.017.

(83) Lumpkin, R. J.; Komives, E. A. DECA, A Comprehensive, Automatic Post-Processing Program for HDX-MS Data. Mol. Cell. Proteomics MCP 2019, 18 (12), 2516–2523. 10.1074/mcp.TIR119.001731.

